# Elevated temperature increases genome-wide selection on de novo mutations

**DOI:** 10.1101/268011

**Authors:** David Berger, Josefine Stångberg, Julian Baur, Richard J. Walters

## Abstract

Adaptation in new environments depends on the amount and type of genetic variation available for evolution, and the efficacy by which natural selection discriminates among this variation to favour the survival of the fittest. However, whether some environments systematically reveal more genetic variation in fitness, or impose stronger selection pressures than others, is typically not known. Here, we apply enzyme kinetic theory to show that rising global temperatures are predicted to intensify natural selection systematically throughout the genome by increasing the effects of DNA sequence variation on protein stability. We tested this prediction by i) estimating temperature-dependent fitness effects of induced random mutations in seed beetles adapted to ancestral or warm temperature, and ii) calculating 100 paired selection estimates on mutations in benign versus stressful environments from a diverse set of unicellular and multicellular organisms. Environmental stress *per se* did not increase the mean strength of selection on de novo mutation, suggesting that the cost of adaptation does not generally increase in new environments to which the organism is maladapted. However, elevated temperature increased the mean strength of selection on genome-wide polymorphism, signified by increases in both mutation load and mutational variance at elevated temperature. The theoretical predictions and empirical data suggest that this increase may correspond to a doubling of genome-wide selection for a predicted 2-4°C climate warming scenario in ectothermic organism living at temperatures close to their thermal optimum. These results have important implications for global patterns of genetic diversity and the rate and repeatability of evolution under climate change.

**Impact Statement:** Natural environments are constantly changing so organisms must also change to persist. Whether they can do so ultimately depends upon the reservoir of raw genetic material available for evolution, and the efficacy by which natural selection discriminates among this variation to favour the survival of the fittest. Here, the biochemical properties of molecules and proteins that underpin the link between genotype and phenotype can exert a major influence over how the physical environment affects the expression of phenotypes and the fitness consequences of DNA sequence polymorphism. Yet, the constraints set by these molecular features are often neglected within eco-evolutionary theory trying to predict evolution in new environments. Here we combine predictions from existing biophysical models of protein folding and enzyme kinetics with experimental data from ectothermic organisms across the tree of life, to show that rising global temperatures are predicted to increase the mean strength of selection on DNA sequence variation in cold-blooded organisms. We also show that environmental stress *per se* generally does not increase the mean strength of selection on new mutations, suggesting that genome-wide natural selection is not stronger in new environments to which an organism is maladapted. Theoretical predictions and data suggest that an expected climate warming scenario of a 2-4°C temperature raise within the forthcoming century will result in roughly a doubling of genome-wide selection for organisms living close to their thermal optima. However, our results also point to substantial variability in the temperature-dependence of selection on different proteins within and between organisms, suggesting scope for compensatory adaptation to shape this relationship. These results bear witness to and extend the universal temperature dependence of biological rates and have important implications for global patterns of genetic diversity and the rate and repeatability of genome evolution under environmental change.

## Introduction

The strength of natural selection impacts on the rate and repeatability of evolution (Haldane 1927; Fisher 1930; Kimura 1968; Orr 2005; Svensson and Berger 2019), the maintenance of genetic variation (Lande 1975; Turelli 1984) and extinction risk (Haldane 1937; Bürger and Lynch 1995). However, despite being central to predicting the impacts of climate change, there is little information about whether some environmental factors impose stronger selection pressures than others (Chevin et al. 2010; Kokko et al. 2017). Mapping of the environment’s influence on phenotype is therefore of paramount importance to understanding species persistence (Kingsolver et al. 2001; Agrawal and Whitlock 2010; Siepielski et al. 2013, 2017; Caruso et al. 2017). It is widely recognized that environmental change should increase the strength of directional selection on the key traits and genes underlying local adaptation. However, the fitness consequences associated with maladaptation at such genes may be relatively small compared to the variance in fitness attributed to segregating polymorphisms across the entire genome (Haldane 1937; Agrawal and Whitlock 2012). This reservoir of genetic variation is expected to have fundamental impact on adaptability and extinction risk (Haldane 1937; Bürger and Lynch 1995; Agrawal and Whitlock 2012), but how the environment influences the expression and consequences of genome-wide polymorphism remains poorly understood (Martin and Lenormand 2006; Agrawal and Whitlock 2010; Chevin et al. 2010; Caruso et al. 2017). For example, it is sometimes argued that fitness effects of sequence variation are magnified in new environments due to compromised phenotypic robustness and the release of “cryptic genetic variation” (de Visser et al. 2003; Paaby and Rockman 2014; Siegal and Leu 2014). Yet, empirical evidence for this prediction remains ambiguous (Hoffmann and Merilä 1999; Rowiński and Rogell 2017; Noble et al. 2019) and others have argued that environmental change will have idiosyncratic effects on the mean strength of selection (Martin and Lenormand 2006; Agrawal and Whitlock 2010). These conflicting predictions suggest that only by understanding the mechanistic basis for how environments mould phenotypic effects of genetic variation will it be possible to predict adaptation under future climate change.

Here we demonstrate how considerations of biophysical constraints on protein function can generate insights about how climate change and regional temperatures affect the strength of selection on DNA sequences. Thermodynamics pose a fundamental constraint on protein function, and although there is room for organismal adaptation to circumvent these deterministic effects (Hochachka and Somero 2002; Clarke 2004; Angilletta 2009), there is a strong temperature-dependence of ectotherm behaviour, life-history and fitness (Huey and Kingsolver 1989; Hochachka and Somero 2002; Brown et al. 2004; Angilletta 2009). By adopting a biophysical model of enzyme kinetics and protein stability, we demonstrate that elevated temperatures are predicted to cause a mean increase in the fitness effects of de novo mutation.

We test these predictions by measuring selection on induced mutations at benign and elevated temperature in experimental evolution lines of the seed beetle, *Callosobruchus maculatus*, adapted to ancestral or warm temperature. Second, we analyse 100 published estimates of paired selection coefficients against de novo mutations in benign versus stressful environments from a diverse set of ectothermic organisms, ranging from unicellular bacteria and fungi, to multicellular plants and animals. Our analyses demonstrate that environmental stress *per se* does not affect the mean strength of selection but provide support for the prediction that elevated temperature leads to an average increase in genome-wide selection and genetic variance in fitness. These results have important implications for extinction risk and global patterns of genetic diversity and suggest that the evolution of DNA sequences could proceed at an ever-accelerating pace under continued climate warming.

## Methods

### Enzyme kinetics theory predicts temperature-dependence of mutational fitness effects

Ectotherm fitness shows a well-characterised empirical relationship with temperature that closely mirrors the thermodynamic performance of a rate-limiting enzyme (Huey and Kingsolver 1989; Hochachka and Somero 2002; Angilletta 2009). This empirical relationship reflects the fact that biological rates are governed at the biochemical level by the enzymatic catalytic rate, *k_cat_*, described by the Eyring equation:

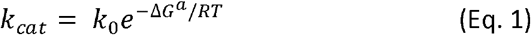

Where *k*_0_ is a rate-specific constant, *ΔG^a^* is the Gibbs free energy of activation required for the enzymatic reaction (kcal mol^−1^), *R* is the universal gas constant (0.002 kcal mol^−1^) and *T* is temperature measured in degrees Kelvin (Schoolfield et al. 1981). Equation (1) thus describes an exponential increase in reaction rate with temperature (Hochachka and Somero 2002).

The observed decline in biological rate at temperatures exceeding the organism’s thermal optimum is attributed to a reduction in the proportion of functional enzyme due to reversable inactivation via protein unfolding (DePristo et al. 2005; Tokuriki and Tawfik 2009; Sikosek and Chan 2014; Bershtein et al. 2017; Echave and Wilke 2017) (Fig. 1a,b). Protein folding can be described by a Boltzmann probability as a function of the Gibbs free energy of folding, *ΔG^f^*, which is thus a measure of protein stability (DePristo et al. 2005):

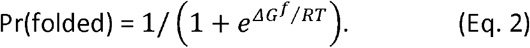

**Figure 1:**
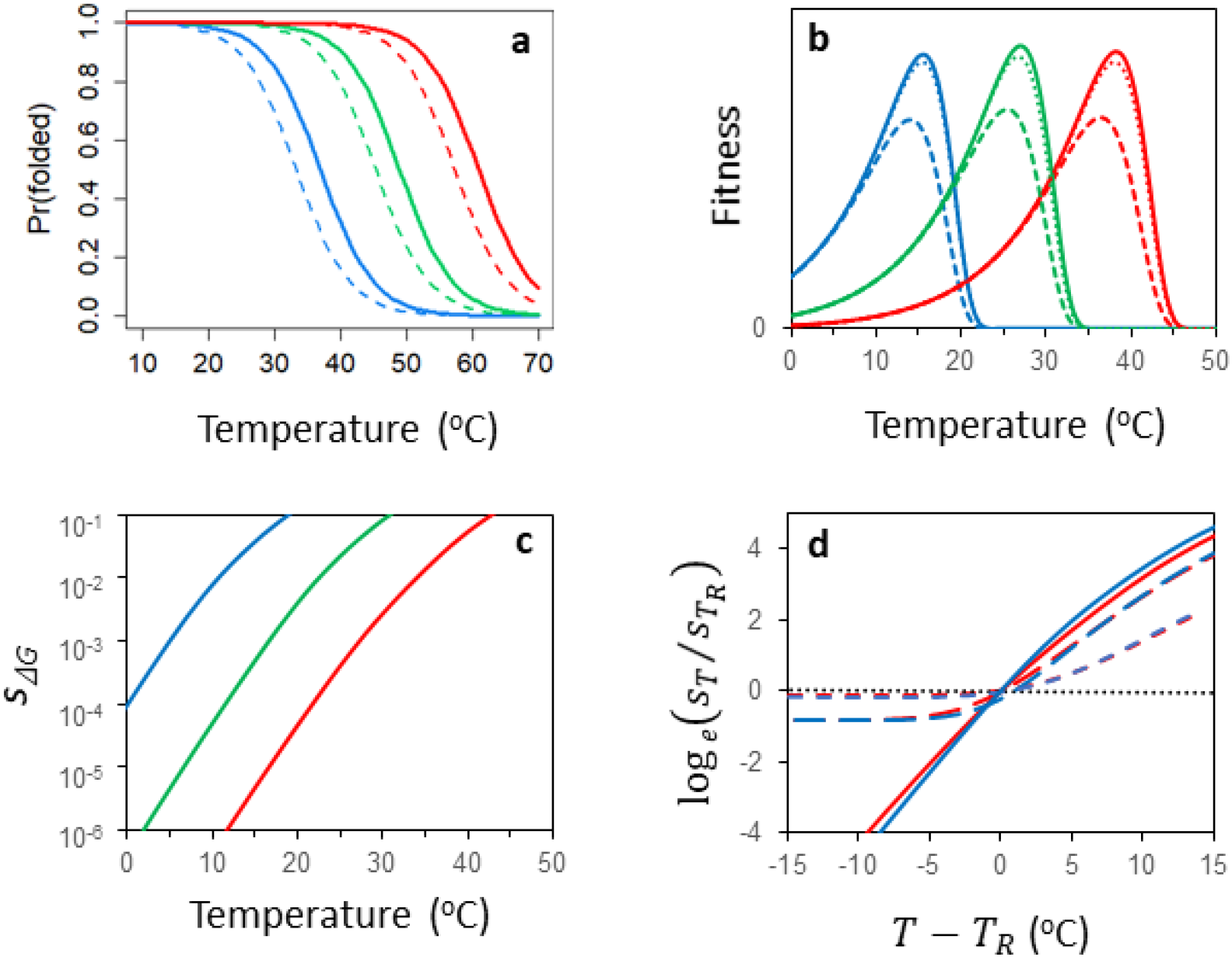
An enzyme-kinetic model of temperature dependent mutational fitness effects. In a) the fraction of folded protein for a wildtype (full lines) and mutant protein with a single stability mutation (*ΔΔ^f^* = 0.9, dashed lines). Illustrated for an unstable (blue) intermediate (green) and stable (red) protein (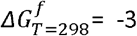, 6 and −9 kcal/mol, respectively). In b) expected fitness for three genotypes with an ensemble of 500 multiplicatively acting proteins with different mean stabilities 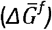 (blue, green and red lines reflect 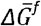 values of −6, −9 and −12 kcal/mol and *ΔG^a^* values = 19.25, 20.00 and 20.76 kcal/mol). Solid lines = wildtype, short-dashed lines = mutant carrying a single folding mutation, long-dashed lines = mutant carrying 10 folding mutations. Mutations were drawn from the empirical normal distribution (*ΔΔG^f^* = 0.9 ± 1.7 SD). In c) the expected mean selection coefficient against a single folding mutation occurring at a random gene for each of the three genotypes. In d, ‘warm’ and ‘cold’ adapted genotypes experience equivalent strengths of selection when fitness effects are assessed at a temperature standardised relative to each genotype’s thermal optimum (*T_R_ − T_OPT_* − 10 °C, for clarity only reaction norms for 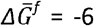 and −12 are shown). Mutational fitness effects on catalytic rate (*ΔΔG^a^*) show no discernible temperature dependence (black dotted line; here *ΔΔG^a^* lowers fitness at *T_opt_* by s = 10^−2^). However, if a mutation has pleiotropic effects on both *ΔΔG^f^* and *ΔΔG^a^*, the temperature dependence of selection against the mutant brought about by its effects on stability can be masked (full, long-dash and short-dash lines equate to a s(*ΔΔG^a^*) = 0, 10^−3^ and 10^−2^ at *T_opt_*). All examples used a value of *ΔS^f^* = 0.25kcal/mol K^−1^ and assumed that mutations affected stability via changes in enthalpy (*ΔΔH^f^*). See main text and Supplementary 1 for further details.

At a benign temperature of 25°C, most natural proteins occur in functional state and the value of the Gibbs energy is negative (mean *ΔG^f^* ^~^ −7 kcal mol^−1^; DePristo et al. 2005; Chen and Shakhnovich 2009). However, both *ΔG^a^* and *ΔG^f^* are comprised of an enthalpy term (Δ*H*) and a temperature-dependent entropy term (Δ*S*) that reduces to: Δ*G*(*T*) = Δ*H* + *T*Δ*S* over the ecologically relevant temperature range of most organisms (Chen and Shakhnovich 2010). Based on values from the literature (Chen and Shakhnovich 2010; Dill et al. 2011), Δ*S^f^* ^~^ 0.25-1 kcal/mol K^−1^ for a protein of typical length. From equation (2) it is therefore clear that warm temperatures increase entropy making *ΔG^f^* less negative, which leads to a rapid and non-linear reduction in the fraction of functional enzyme. This non-linearity suggests that any mutational change in protein stability should be more consequential for folding when temperatures rise (Fig 1a).

The reaction rate kinetics of equation (1) can be combined with the protein folding of equation (2) to describe the “fitness” of a rate-determining protein, *i*, as a function of temperature and the Gibbs energies for activation and folding (Chen and Shakhnovich 2010) (Fig. 1b):

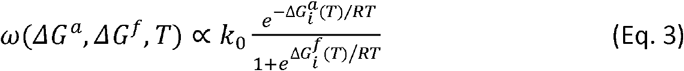

Here we use equation (3) to explore the effect of temperature on the strength of selection on de novo mutations that impact catalytic rate and/or protein folding stability. We introduce a mutational change in the Gibbs free energy of activation (*ΔΔG^a^*) and folding (*ΔΔG^f^*), respectively:

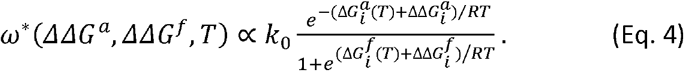

We then calculate mean selection against a de novo mutation:

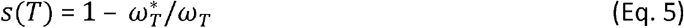

where ***ω***_T_* and ***ω***_T_ is fitness of the mutant and the wildtype at temperature *T*. Assuming fitness effects are multiplicative across proteins, substituting equations (3) and (4) into equation (5) yields the following expression:

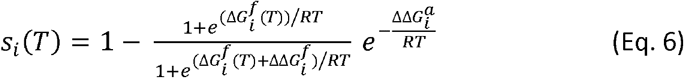

Based on equation (6), selection against mutations affecting enzymatic reaction rate (*ΔΔG^a^*) is expected to be independent of the original catalytic rate of the wildtype (*ΔG^a^*) and remain largely unaffected by any ecologically relevant (i.e. 283-313°K) change in temperature (Fig. 1c, see further: Supplementary 1.1). Note, however, from equations (2) and (4) and Fig. 1a how mutation and temperature-driven increases in entropy both have non-linear effects on protein folding. To illustrate the consequences of this synergism we applied equation (6) in numerical simulations to calculate the expected mean selection on a mutation in a random protein with its original stability drawn from a truncated gamma distribution (*ΔG^f^* ^~^ − Γ(*k* = 5.50, *θ* = 1.89) derived from empirical data from bacteria, yeast and nematodes (Ghosh and Dill 2010). We compared the resulting temperature dependence of selection in three enzyme ensembles with different hypothetical distributions of protein stabilities by shifting the empirical gamma distribution so that mean *ΔG^f^* = −6, −9 or −12, respectively (Fig. 1a-d). In the example presented in figure 1, we set *ΔS^f^* = 0.25, but in Supplementary 1 we show results for different values of *ΔS^f^*.

The majority of mutations with effects on folding (*ΔΔG^f^*) are expected to destabilize proteins (DePristo et al. 2005; Drummond and Wilke 2008; Sikosek and Chan 2014; Echave and Wilke 2017) and the net impact of a single mutation has been estimated to *ΔΔG^f^* ^~^ 0.9 kcal mol^−1^ (SD = 1.7) (Zeldovich et al. 2007; Tokuriki and Tawfik 2009), a value more or less independent of the original protein stability (*ΔG^f^*) (Chen and Shakhnovich 2009). Hence, each protein was mutated by sampling a single folding mutation from this empirically estimated normal distribution. Empirical estimates of mutational effects on the activation energy of random enzymes are likely very biased (see Supplementary 1.1). In the simulations we therefore chose parameter values of *ΔΔG^a^* that yielded negative selection coefficients of magnitudes that are typically observed at benign temperatures (*s* = 0–10^−2^, T=298°K, see Supplementary 1.1). Mutational effects on the Gibbs energy (*ΔΔG*) can derive from a combination of both enthalpic (*ΔΔH*) and entropic (*ΔΔS*) terms. In the examples in Figure 1 we assumed that all mutational effects are enthalpic in origin. However, qualitative conclusions hold when mutational effects are modelled through only the entropic term or enthalpic and entropic terms combined (Supplementary 1.2). R code available upon request. We centre our discussion in the *Results* on the temperature-dependence of the mean strength of selection, as these predictions were most readily testable with our collated empirical data (see further below). However, the temperature-dependence of selection on individual mutations is predicted to be variable both across genes and organisms, and more quantitative predictions strongly depend on model assumptions, which is further explored in Supplementary 1.

### Temperature-dependent fitness effects of mutations in seed beetles adapted to contrasting thermal regimes

To test model predictions, we measured fitness effects of induced random mutations at 30⍰C and 36⍰C in lines of the seed beetle *Callosobruchus maculatus*, evolved at benign 30⍰C (three ancestral lines) or stressful 36⍰C (three warm-adapted lines) for more than 70 generations (see Supplementary 2). Previous studies have revealed significant differentiation in key life history traits between the regimes (Rogell et al. 2014; Berger et al. 2017). Here we further detailed thermal adaptation across the regimes by first analysing differences in offspring production in the lines at the two assay temperatures in the current experiment (see further below). Second, we quantified reaction norms for juvenile survival and development rate across five temperatures (23, 29, 35, 36 & 38°C) following 100 generations of experimental evolution and two generations of common-garden acclimation at 30°C to remove non-genetic parental effects. Newly laid eggs were subsequently randomized to each assay temperature with larval food provided ad libitum. Effects of assay temperature and selection regime on survival and development rate (1/development time) of 2755 offspring, evenly split over the five assay temperatures and six replicate lines, were analysed in linear mixed effects models in the lme4 package (Bates et al. 2014) for R (see further: *Extended Supplementary Methods*).

We compared fitness effects of induced mutations at 30°C and 36°C for each line of the two evolution regimes. A graphical depiction of the design can be found in Supplementary 2. All six lines were maintained at 36⍰C for two generations of common-garden acclimation. The emerging virgin adult offspring of the second generation were used as the F0 individuals of the experiment. We induced mutations by exposing half of the F0 males to gamma radiation at a dose of 20 Grey (20 min treatment), with the other half kept as controls. All males were mated with virgin females from their own population at benign 30°C. The mated females were immediately placed on beans presented ad libitum and randomized to a climate cabinet set to either 30°C or 36°C (50% RH) and allowed to lay their lifetime storage of F1 eggs. To exclude non-genetic effects of the irradiation treatment, we measure mutational effects in the F2 generation. To this end we applied a Middle Class Neighborhood breeding design to nullify selection on all but the unconditionally lethal mutations amongst F1 juveniles (Shabalina et al. 1997); from the F1 survivors, we crossed two randomly selected male and female offspring per family with another family from the same treatment and line, so that each family contributed equally to the F2 generation (irrespective of the mutation load it carried). This approach allowed us to quantify the cumulative deleterious fitness effect of all but the unconditionally lethal mutations induced in F0 males (i.e. mutation load) by comparing the production of F2 adults in irradiated lineages, relative to the number of adults descending from F0 controls (Fig. S2). We also used F1 adult counts to derive an estimate of load, acknowledging that it may include non-trivial paternal effects from the irradiation treatment, in addition to pure mutational effects. However, results based on F1 and F2 estimates were consistent (Fig 3). To estimate the effects of elevated temperature on mutational fitness effects in the two genetic backgrounds, we analysed the number of offspring produced as a Poisson response, using generalized linear mixed effects models, testing for interactions between radiation treatment, assay temperature and evolution regime. To better illustrate the results graphically, we also ran Bayesian analyses using the MCMCglmm package (Hadfield 2010) with the same model structure, but assuming a normally distributed response. We then calculated the posterior estimates of mutation load in each environment for each evolution regime as: **Δ*ω* = 1- *ω*_IRR_/ω_CTRL_** (where ***ω*** is the number of adult offspring) directly from these models (Fig. 3) (see *Extended Supplementary Methods*).

### A meta-analysis of temperature dependent mutational fitness effects

To test the hypothesis that elevated temperature increases the mean strength of genome-wide selection more generally, we retrieved 100 paired estimates comparing selection on de novo mutations across a benign and stressful environment from 28 studies on 11 organisms, spanning viruses and unicellular bacteria and fungi, to multicellular plants and animals. The original studies measured effects of mutations accrued by mutation accumulation, mutagenesis, or targeted insertions/deletions. This was done by estimating fitness in form of Malthusian growth rate, survival, or reproduction in mutants relative to wildtypes (Supplementary 3). Hence, the strength of selection against mutants in each environment (i.e. environment-specific mutation load) could be estimated as: Δ***ω**_i_* = 1-***ω**_i_^*^*/***ω**_i_*, where ***ω**_i_^*^* and ***ω**_i_* is the fitness in environment *i* of the mutant and wildtype respectively. An estimate controlling for between-study variation was retrieved by taking the log-ratio of mutation load in the stressful relative to benign environment in each study: Log_e_[***Δω***_stress_/**Δ*ω***_benign_], with a log-ratio above (below) 0 indicating stronger (weaker) selection against mutations under environmental stress.

We analysed the log-ratios in meta-analysis using Bayesian mixed effects models to estimate if log-ratios differed from 0 for three levels of environmental stress: cold temperature, warm temperature, and other types of stress pooled, as well as for the total effect of stress averaged across all studies. We also tested if log-ratios differed between the three types of abiotic stress. Each estimated log-ratio’s contribution to the final meta analytic result was weighted by its sampling variance by passing the standard error (SE) of each log-ratio to MCMCglmm using the idh(SE):units command. The standard errors were calculated by propagation of measurement errors from the original studies. All models included stress-type, mutation induction protocol and fitness estimate as main effects, although effects of the latter two were never significant. We included study organism and study ID as random effects. Additionally, study organism was crossed with stress type to control for species variation and phylogenetic signal. To further explore large scale signals in the data we performed an analysis including a fixed factor encoding uni- or multicellularity, which was crossed with stress type, allowing us to test for differences in selection between the two groups (see *Supplementary Methods*). We explored any potential publication bias by plotting the precision of each estimate (1/standard error) against its mean. This showed no clear evidence for such bias (Supplementary 3).

## Results

### Enzyme kinetics theory predicts temperature-dependence of mutational fitness effects

Enzyme kinetic theory parameterized with empirical estimates of protein stabilities (Eq. 6) predicts that the mean strength of selection against random de novo mutations should increase with temperature (Fig. 1c). This effect is further attributed to the synergistic effect of temperature and de novo mutation to destabilize proteins (Fig. 1a). In Supplementary 1.2 and 1.3 we show that this is also predicted to lead to a substantial increase in evolutionary potential in form of mutational variance in fitness and a larger fraction of both highly deleterious and beneficial mutations. Second, the evolution of increased protein thermostability is predicted to produce enzymes that are more robust to mutational perturbation (Fig. 1a, c), and hence, mutations in proteins with low stability are predicted to contribute disproportionally to temperature dependent mutational fitness effects (Supplementary 1.3, see also: Agozzino and Dill 2018). Third, if differences in thermostability of proteins evolve in response to environmental temperatures, cold- and warm-adapted ecotypes are predicted to experience the same strength of selection on de novo mutations at an environmental temperature standardized relative to the ecotypes’ respective thermal optima (Fig. 1d). We note here, however, that while individual proteins isolated from mesophilic and thermophilic organisms do show evolved differences in stabilities, this seem not to hold for the organismal-wide repertoires of proteins measured in vitro (Leuenberger et al. 2017). Fourth, variation in the temperature dependence of mutational effects might be generated by differences in *ΔS^f^*, governing the temperature sensitivity of protein stability (i.e. *ΔG^f^*(*T*)) (Supplementary 1.4, see also Sawle and Ghosh 2011). Hence, this could account for variability in the temperature-dependence of selection across genes within an organism, or general variability in the temperature-dependence of selection between organisms. Fifth, while the mean mutational effect on catalytic rate (*ΔΔG^a^*) is predicted to be largely independent of temperature (Supplementary 1.1), *ΔΔG^a^* mutations are expected to weaken the overall temperature-dependence of mutational fitness effects. The extent to which they do depends on their effect size relative to that of folding mutations (*ΔΔ^f^*) (Fig. 1d).

These predictions arise from two well-established principles: i) enzymes show disproportionate reversible inactivation at high temperatures driven by increases in the entropy of folding (Hochachka and Somero 2002), and ii) the majority of de novo mutations act to destabilize proteins (DePristo et al. 2005; Drummond and Wilke 2008; Tokuriki and Tawfik 2009; Sikosek and Chan 2014; Bershtein et al. 2017; Echave and Wilke 2017). The qualitative results are therefore robust to the particular mathematical formulation of the enzyme-kinetic model. In Supplementary 1.5 we also show that a model of metabolic flux theory incorporating protein toxicity (Serohijos and Shakhnovich 2014; Echave and Wilke 2017) makes the same qualitative predictions as outlined above. This model makes the additional prediction that the temperature dependence should, all else equal, be stronger for more abundant proteins. This is in line with documented greater fitness costs of folding mutations in more expressed genes, which has been postulated as a reason for why such genes evolve more slowly (Drummond et al. 2005; Serohijos et al. 2012).

### Temperature-dependent fitness effects of mutations in seed beetles adapted to contrasting thermal regimes

As expected, there was considerable signs of thermal adaptation across the compared evolution regimes of seed beetle; offspring production assayed in the non-irradiated lineages kept at the two temperatures (30⍰C vs 36⍰C) demonstrated local adaptation to temperature (Fig. 2a). Thermal reaction norms for juvenile development rate and survival showed that warm-adapted lines develop slightly slower but survive slightly better across the temperature range (Fig. 2b-c), a sign of counter-gradient adaptation (Hochachka and Somero 2002; Angilletta 2009) (i.e. counteracting the general effect of elevated temperature to increase development rate and decrease survival).

**Figure 2:**
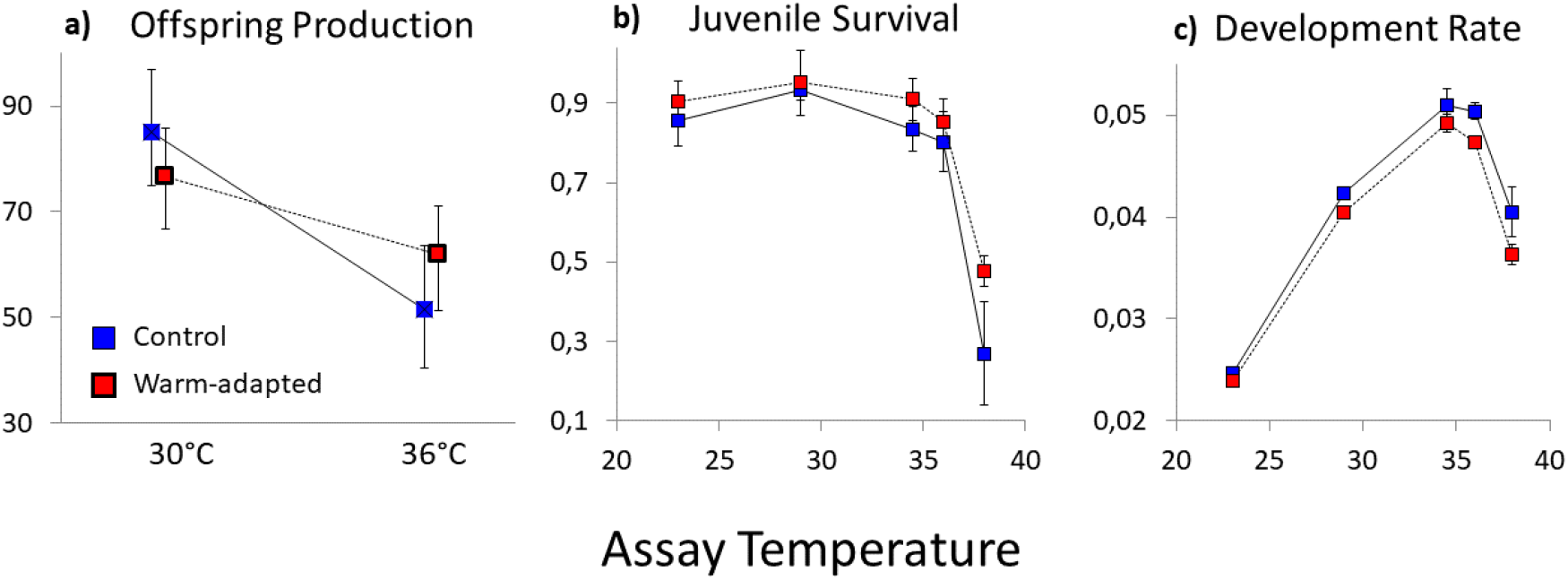
Thermal adaptation during experimental evolution. The level of adaptation to simulated climate warming measured as (A) adult offspring production at 30 and 36°C, and thermal reaction norms for (B) juvenile survival and (C) development rate (means ± 95% confidence limits). Blue and red symbols denote control and warm-adapted lines, respectively. While offspring production was generally decreased at 36⍰C relative to 30⍰C (X^2^ = 62.5, df = 1, P < 0.001, total n = 698), this decrease was less pronounced in warm-adapted lines (interaction: X^2^ = 7.35, df = 1, P = 0.007; Fig. 2a) signifying significant local adaptation to respective thermal regime. Warm-adapted lines showed consistently slower development (X^2^ = 27.2, df= 1, P < 0.001, total n = 2755, Fig 2b) and marginally higher survival (X^2^= 3.74, df= 1, P = 0.053, total n = 2755, Fig. 2c) across temperatures, in line with counter-gradient adaptation in response to the effect of elevated temperature to increase juvenile development rate but decrease juvenile survival.

Despite the observed thermal adaptation, there was no obvious difference in the temperature-dependence of mutational fitness effects seen in control and warm-adapted lines (interaction: P_F1_ = 0.43, P_F2_ = 0.90; Fig. 3); Elevated temperature caused a general increase in mutation load in both the F1 (X^2^ = 13.0, df = 1, P < 0.001, total n = 713, Fig. 3a) and F2 generation (X^2^ = 7.44, df = 1, P = 0.006, total n = 1449, Fig 3b), consistent with model predictions (compare Fig. 1c and Fig. 3). The fact that mutational fitness effects in control and warm-adapted genotypes, locally adapted to the alternative assay temperatures, showed qualitatively similar thermal dependence suggests that high temperature, rather than stress *per se*, caused the increase in selection. We note here that, while the greater overall mutational effects seen in control relative to warm adapted lineages also are consistent with model predictions (compare Fig. 1c and Fig 3), this difference is not easily interpretable and may have several alternative explanations (see *Extended Supplementary Methods*).

**Figure 3:**
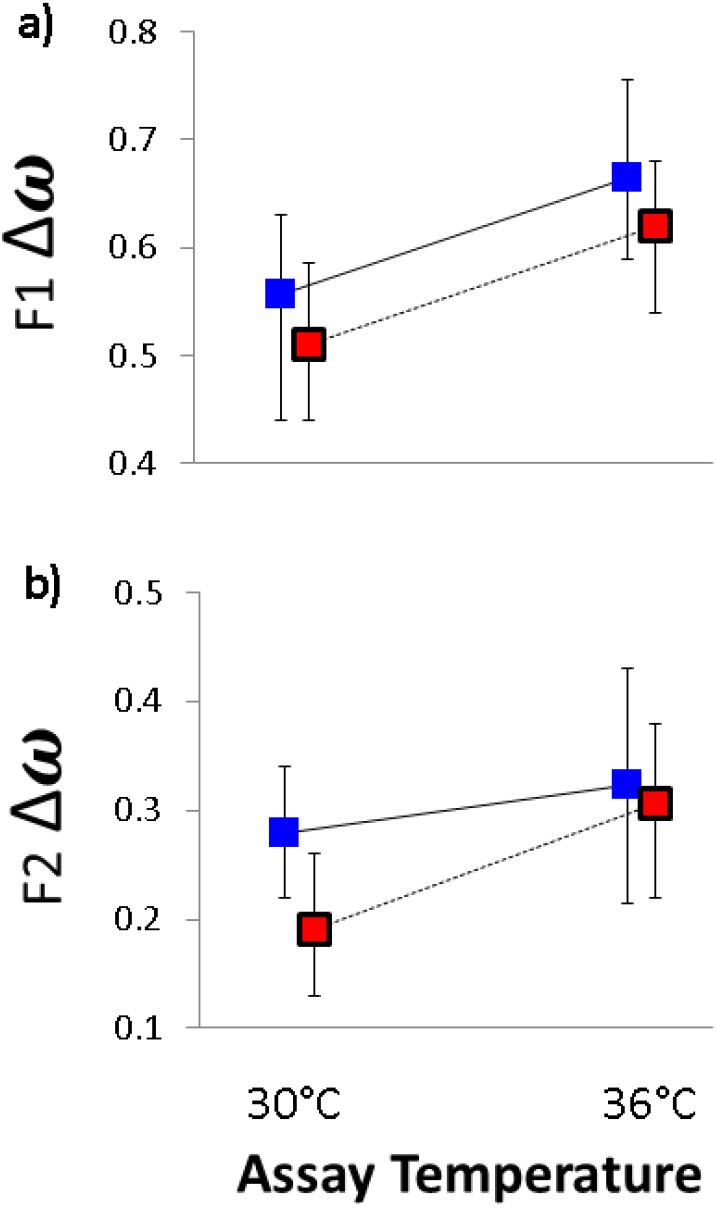
The evolution of temperature dependent mutational fitness effects. Mutation load (**Δ*ω***) (mean ± 95% confidence limits) measured for (A) F1 juvenile survival and (B) F2 adult offspring production, at the two assay temperatures. There was an overall strong and significant increase in **Δ*ω*** at hot temperature. This effect was similar across the three control (blue) and three warm-adapted (red) lines, in both the F1 and F2 generation.

### A meta-analysis of temperature dependent mutational fitness effects

Analysing all collated log-ratios together showed that selection against de novo mutation was not greater under stressful abiotic conditions on average (log-ratio = 0.19, 95% CI: −0.07-0.45; P_MCMC_ = 0.13, Fig 4). We continued by analysing the 40 estimates derived at high and low temperature stress separately from the 60 estimates derived from various other stressful environments (summarized in Supplementary 3). This revealed that selection on de novo mutation increases at elevated temperature (log-ratio ≤ 0; P_MCMC_ < 0.001, n = 21, studies = 10), whereas there was no increase at low temperature (log-ratio ≤ 0; P_MCMC_ = 0.67, n = 19, studies = 11) or for the other forms of stress pooled (log-ratio ≤ 0; P_MCMC_ = 0.48, n = 54, removing 6 estimates that were not possible to analyse due to *s* in each environment ^~^ 0, studies = 22). Elevated temperature led to a significantly larger increase in selection relative to both cold stress (P_MCMC_ = 0.004) and the other stressors pooled (P_MCMC_ = 0.002) (Fig 4). There was a tendency for cold stress to decrease selection in unicellular species and increase it in multicellular species, but this effect was marginally non-significant (interaction: P_MCMC_ = 0.066). Moreover, 5 of the 6 estimates at cold stress for multicellular species derive from survival data on *D. melanogaster* and drive this trend (Fig. 5b). We found no evidence for differences in the effect of elevated temperature on selection between the four multicellular and three unicellular species in the data (P_MCMC_ = 0.45); mutational effects were greater at elevated temperature in 8/10 and 10/11 cases in multicellular and unicellular species, respectively (combined two-sided binomial test, P = 0.0015). Notably, the 12 log-ratios that were significantly different from 0 (>1.96SE) at elevated temperature were all positive, signifying increased selection (two-sided binomial test, P = 0.0005). An analysis of 64 paired estimates of mutational variance followed the same general pattern as mutation load (Fig. 4 and Supplementary 3), as predicted by the biophysical model (Supplementary 1.2 and 1.3).

**Figure 4:**
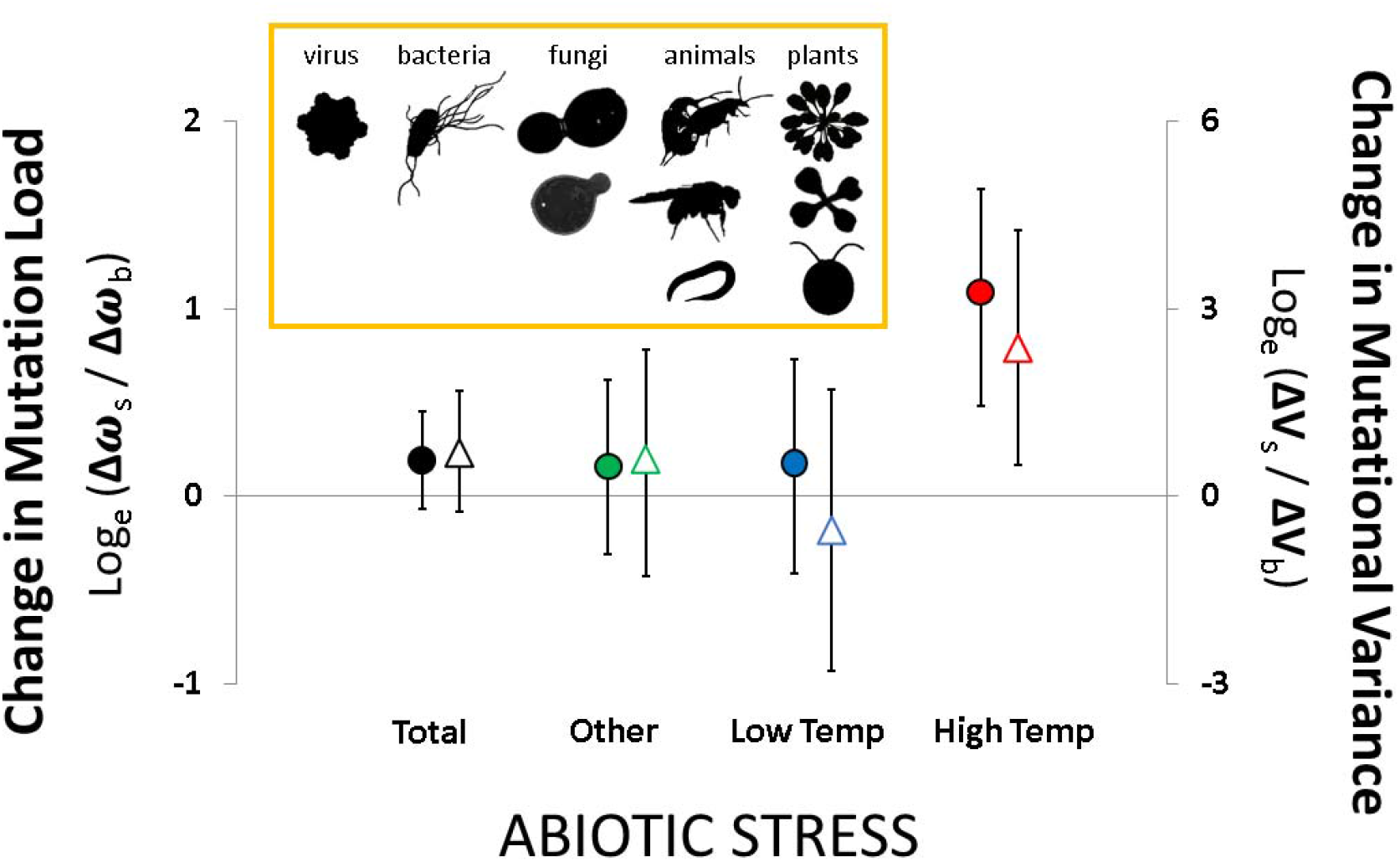
Meta-analysis of mutational fitness effects in stressful environments. Meta-analysis of the effect of abiotic stress on the mean strength of selection against de novo mutations (filled points) and mutational variance (open triangles) analysed by log-ratios (Bayesian posterior modes ± 95% credible intervals): **Δ** _stress_/ _benign_ and **ΔV**_stress_/ _benign_ > 0 correspond to greater mutational fitness effects under environmental stress. The 94 paired estimates of **Δ** (filled circles) show that selection is not greater in stressful environments overall (P_MCMC_ = 0.13) and highly variable across the 25 studies analysed. However, estimates of **Δ** at high temperature are greater than their paired estimates at benign temperature (P_MCMC_ < 0.001). Results were similar when analysing the fewer available estimates of mutational variance (**ΔV**: open triangles, P_MCMC_ = 0.02). The box shows the eleven species included in the analysis (of which two were roundworms), covering four major groups of the tree of life. See main text and Supplementary 3 for further details.

**Figure 5:**
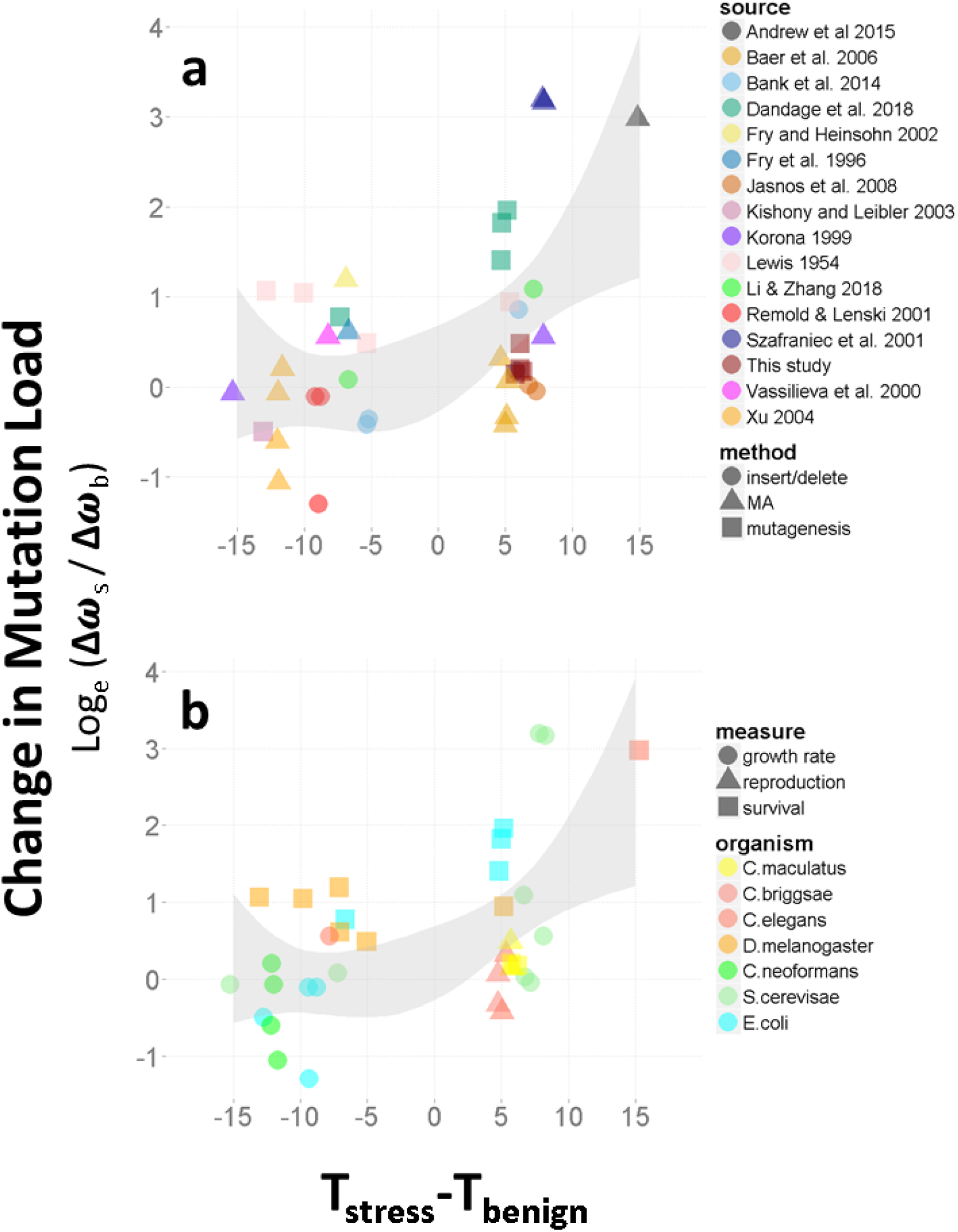
Meta-analysis of temperature-dependent mutational fitness effects. Temperature-dependent mutational fitness effects. The strength of selection on de novo mutations as a function of the direction and magnitude of the temperature shift between the benign and stressful temperature. In (A) the 16 studies analysed and the method used to induce mutations, is depicted. In (B) the seven species analysed, and the fitness measure taken, is depicted. Selection generally increases with temperature (P_MCMC_ < 0.001) whereas stress per se (quantified as the mean reduction in relative fitness between the benign and stressful temperature) did not affect the strength of selection (P_MCMC_ > 0.8). The grey shaded area represents the 95% CI from a second degree polynomial fit of the log-ratios on temperature, weighted by the statistical significance of each estimate (absolute log-ratio/standard error). Points are jittered for illustrative purposes.

Using the 40 paired estimates of mutation load at contrasting temperatures we partitioned effects on the strength of selection from i) stress *per se*; quantified as the reduction in mean fitness at the stressful temperature relative to the benign temperature 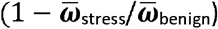, and ii) that of the temperature shift itself; quantified as the magnitude and direction of the temperature shift: T_stress_ – T_benign_. The strength of selection was not related to stress (P_MCMC_ > 0.8). However, a shift towards warmer assay temperature *per se* caused a substantial increase in mutation load (P_MCMC_ < 0.001, Fig 5). There was also a non-linear effect of temperature (P_MCMC_ = 0.01, Fig 5). Again, the effect of cold temperature on the strength of selection seemed to differ between the unicellular and multicellular species studied (interaction: P_MCMC_ = 0.018, Fig. 5b). However, given that this pattern is driven almost solely by the 5 estimates from *D. melanogaster*, more data is needed to say anything concrete about this potential effect of cellularity at cold temperature.

The long term consequences of the revealed relationship will depend on if the predicted effects of temperature on protein stability will change the relative abundance of nearly neutral to strongly deleterious alleles (Kimura 1983; Bershtein et al. 2006; Agrawal and Whitlock 2012; Siegal and Leu 2014). In Supplementary 3 we show that increases in both the number of (conditionally) expressed mutations as well as increases in their average fitness effect are likely to underlie the observed increases in Δ***ω*** at elevated temperature.

## DISCUSSION

Here we have presented evidence suggesting that elevated temperature increases the mean strength of genome-wide selection and mutational variance in fitness, an observation qualitatively consistent with the applied biophysical model of enzyme kinetics, which ascribes these increases to magnified allelic effects on protein folding at elevated temperature. In contrast, environmental stress *per se* did not have an effect on the strength of selection in any analysis, implying that mutational robustness is not compromised in stressful environments (Martin and Lenormand 2006; Agrawal and Whitlock 2010). The empirical data suggest that, without adaptation, the depicted scenario of 2-4°C of warming by the end of this century (Fifth Assessment Report - Climate Change 2013) could result in a doubling of genome-wide selection on average. However, the existing data at hand also hint at there being variation between organismal groups in the effect (Fig. 5), a possibility that needs further empirical exploration. Indeed, the exact strength of the temperature dependence of selection is predicted to vary between genes and mutations and depends on model assumptions (Fig. 1, Supplementary 1). Moreover, even though there is a strong qualitative agreement between empirical data and the theoretical predictions, further in-depth studies are urgently needed to prove causality and evaluate model assumptions in more detail.

An increase in mutational effects at warm temperature is predicted to influence regional patterns of evolutionary potential. Previous studies have highlighted a range of possible consequences of temperature on evolutionary potential in tropical versus temperature regions (Allen et al. 2002, 2006; Mayhew et al. 2008, 2012; Tittensor et al. 2010; Puurtinen et al. 2016; Yasuhara and Danovaro 2016), including faster generation times (Gillooly et al. 2002), higher maximal growth rates (Frazier et al. 2006; Walters et al. 2012), higher mutation rates (Berger et al. 2017) and more frequent recombination (Lloyd et al. 2018) in the former. Our results imply that also the efficacy of selection on DNA sequences may be greater in the tropics, which together with the aforementioned factors could result in more rapid evolution and diversification, in line with the greater levels of biodiversity in this area (Jablonski et al. 2006; Tittensor et al. 2010). However, implications for species persistence under climate change will crucially depend on demographic parameters (Bürger and Lynch 1995; Kokko et al. 2017), and given that most mutations are deleterious and act to destabilize proteins, greater purifying selection in tropical areas may result in increased genetic loads and extinction rates if evolutionary potential is limited (Kellermann et al. 2009; Hoffmann and Sgrò 2011; Walters et al. 2012). Moreover, protein stability has itself been suggested to increase evolvability by allowing destabilizing mutations with conditionally beneficial effects on other aspects of protein fitness to be positively selected (Bloom et al. 2006; Wagner 2008, 2017; Soskine and Tawfik 2010). Hence, the destabilizing effect of rising global temperatures on protein folding could by reducing this buffering capacity also limit the potential for evolutionary innovation.

The observed temperature dependence builds a scenario in which climate warming may lead to molecular signatures of increased purifying selection and genome-wide convergence in taxa experiencing climate warming. In support of this claim, Sabath et al. (2013) showed that growth temperatures across thermophilic bacteria tend to be negatively correlated to the non-synonymous to synonymous (dN/dS) nucleotide substitution-rate. Effects could possibly extend to other aspects of genome architecture. For example, Drake (2009) showed that two thermophilic microbes have lower mutation rates than their seven mesophilic relatives and suggested that increased fitness consequences of mutation at hot temperature selects for decreased mutation rate. Increased selection in warm climates could also select for increased mutational robustness (Wagner 2005; Rajon and Masel 2011) via additional mechanisms not explicitly considered by our model, for example via upregulation of chaperones such as heat-shock proteins, known to assist both protein folding and DNA repair (Hochachka and Somero 2002). Indeed, some of the variation in the revealed temperature dependence of mutational fitness effects may be rooted in the use and extent of such compensatory mechanisms (Clarke 2004; Sikosek and Chan 2014; Fields et al. 2015).

Environmental tolerance has classically been conceptualized by a Gaussian function mapping organismal fitness to an environmental gradient (e.g: Levins 1968; Huey and Kingsolver 1989; Bürger and Lynch 1995). In this framework stress is not expected to increase mean selection against de novo mutation (Martin and Lenormand 2006), a prediction supported by our estimates of selection under forms of environmental stress other than temperature (Fig. 4). This framework assumes that phenotypic effects of mutations remain constant. The applied biophysical model differs from this assumption in that mutation and temperature are assumed to have synergistic effects on protein stability, causing mutational effects on protein folding to increase with temperature. While supported by a number of targeted studies on mutant proteins (DePristo et al. 2005; Tokuriki and Tawfik 2009; Sikosek and Chan 2014; Bershtein et al. 2017; Echave and Wilke 2017; Dandage et al. 2018), it remains less clear how these effects map to the level of physiological, morphological and life history phenotypes (Hoffmann and Merilä 1999; Clarke 2004; Berger David et al. 2011; Husby et al. 2011; Berger et al. 2013; Paaby and Rockman 2014; Rowiński and Rogell 2017).

Another open question is how the unveiled temperature-dependence interacts with other features expected to influence the distribution of fitness effects of segregating genetic variants, such as niche width (Chevin et al. 2010; Walters et al. 2012), genome size (Lynch 2007), phenotypic complexity (Orr 2000) and population size (Kokko et al. 2017). Unicellular and multicellular organisms differ in these aspects, and interestingly, our data hint at a difference in mutational fitness effects between the two groups at cold temperature. The model also predicts that the temperature dependence should be weakened by an increased fraction of unconditionally deleterious mutations. Differences between unicellular and multicellular organism could therefore also arise if the link between fitness and rate-dependent processes at the level of enzymes is more direct in unicellular compared to multicellular organisms, resulting in a higher fraction of unconditional mutations and weaker temperature dependence in the latter. Questions such as these will be crucial to answer to understand how the unveiled temperature dependence of mutational fitness effects will affect regional and taxonomic patterns of genetic diversity and evolutionary potentials under climate change.

## Acknowledgements

We thank C. Rüffler, R. Dandage, A. Husby, B. Rogell for valuable input on earlier drafts. We are also grateful to B. Stenerlöw for providing access to the radiation source, and to L. Hallsson, M. Björklund and AA Maklakov for passing on the beetle lines. This work was supported by grants 2015-05223 and 2019-05024 from the Swedish Research Council (VR) to DB.

## Author Contributions

DB performed the experiments on seed beetles together with JS. RJW and DB performed the modelling and DB and JB performed the meta-analysis. DB wrote the manuscript with considerable input from RJW. All authors commented on manuscript drafts.

## Data accessibility

http://dx.doi.org/10.5061/dryad.6dd04

