## Supplementary Methods for "Elevated temperature increases genome-wide selection on de novo mutations"

***Extended Supplementary Methods*:**

**Temperature-dependent fitness effects of de novo mutations in seed beetles**

*Study Populations*

*Callosobruchus maculatus* is a cosmopolitan capital breeder. Adult beetles do not require food or water to reproduce at high rates, starting from the day of adult eclosion (Fox 1993). The juvenile phase is completed in approximately three weeks, and egg to adult survival is above 90% at benign 30°C (Martinossi-Allibert et al. 2017). The lines were derived from an outbred population created by mixing beetles collected at three nearby sites in Nigeria (Fricke and Arnqvist 2007). This population was reared at 30°C on black eyed beans (*Vigna unguiculata*), and maintained at large population size for >90 generations prior to experimental evolution. Replicate lines were kept at 30ᵒC (control lines) or exposed to gradually increasing temperatures from 30°C to stressful 36ᵒC for 20 generations (i.e. 0.3°C/generation) and then kept at 36ᵒC (warm-adapted lines). Population size was kept at 200 individuals for the first 20 generations and then increased to 500 individuals in each line. In this study we compared three replicate lines of each regime.

*Thermal reaction norms for juvenile survival and development rate*

Previous studies have revealed significant differentiation in key life history traits between the regimes (Rogell et al. 2014; Berger et al. 2017). Here we first analysed differences in offspring production in the lines when used in the current experiment at the two assay temperatures (with no mutations induced, see: *Temperature dependent mutational fitness effects* further below). Second, we quantified reaction norms for juvenile survival and development rate across five temperatures (23, 29, 35, 36 & 38°C) following 100 generations of experimental evolution. Two generations prior to the assaying all six lines were moved to 30°C, which is a beneficial temperature to both sets of lines (Fig. 2) (Berger et al. 2017), to ascertain that differences between evolution regimes were due to genetic effects. Newly emerged second generation adults were allowed to mate and lay eggs for 24h on new *V. unguiculata* seeds that were subsequently randomized to each assay temperature in 90mm diameter petri-dishes with ca. 100 seeds per dish with each carrying no more than 4 eggs to make sure larval food was provided ad libitum. Two dishes were set up per temperature for each line. In total we scored egg-to-adult survival and development time for 2755 offspring evenly split over the five assay temperatures and six replicate lines. Survival was analysed using dead/alive as the binomial response, and development rate (1/development time) as a normally distributed response using generalized and general linear mixed effects models, respectively, in the lme4 package (Bates et al. 2014) for R. Temperature and selection regime as well as their interaction were included as fixed effects, and line identity crossed by assay temperature was added as random effect.

*Temperature dependent mutational fitness effects*

We compared fitness effects of induced mutations at 30°C and 36°C for each line of the two evolution regimes. At the onset of our experiments in 2015 and 2016, the populations had been maintained for 70 and 85 generations, respectively. A graphical depiction of the design can be found in Supplementary 2. All six lines were maintained at 36ᵒC for two generations of acclimation. The emerging virgin adult offspring of the second generation were used as the F0 individuals of the experiment.

We induced mutations by exposing the F0 males to gamma radiation at a dose of 20 Grey (20 min treatment). Gamma radiation causes double and single stranded breaks in the DNA, which in turn induces DNA repair mechanisms (Friedberg et al. 2005). Such breaks occur naturally during recombination, and in yeast to humans alike, point mutations arise due to errors during their repair (Friedberg et al. 2005). Newly emerged (0-24h old) virgin males were isolated into 0.3ml ventilated Eppendorf tubes and randomly assigned to either be placed inside a Gamma Cell-40 radiation source (irradiated), or on top of the machine for the endurance of the treatment (non-irradiated). After two hours at room temperature post-irradiation males were emptied of ejaculate and mature sperm by mating with females (that later were discarded) on heating plates kept at 30°C. The males were subsequently moved back to the climate cabinet to mature a new ejaculate. This procedure discarded the first ejaculate that will have contained damaged seminal fluid proteins in the irradiated males (Daly 2012), causing unwanted paternal effects in offspring. Irradiation did not have a mean effect on male longevity in this experiment, nor did it affect the relative ranking in male longevity among the studied populations (Berger et al. 2017), suggesting that paternal effects owing to the irradiation treatment (other than the mutations carried in the sperm) were small. After another 24h, males were mated with virgin females from their own population. The mated females were immediately placed on beans presented ad libitum and randomized to a climate cabinet set to either 30°C or 36°C (50% RH) and allowed to lay their lifetime storage of F1 eggs. We set up 19-38 F0 males (and mating couples) per treatment, assay temperature and line, and 713 males in total.

To measure mutational effects in the F2 generation, we applied a Middle Class Neighborhood breeding design to nullify selection on all but the unconditionally lethal mutations amongst F1 juveniles (Shabalina et al. 1997); from the F1 survivors, we crossed a randomly selected male and female offspring per family with another family from the same treatment and line. From a few treatment:line combinations with a low number of F0 families set up, we did this procedure twice to get a more balanced sample size. This approach allowed us to quantify the cumulative deleterious fitness effect of all but the unconditionally lethal mutations induced in F0 males (i.e. mutation load) by comparing the production of F2 adults in irradiated lineages, relative to the number of adults descending from F0 controls (Fig. S2). We also used F1 adult counts to derive this estimate, acknowledging that it may include non-trivial paternal effects from the irradiation treatment, in addition to pure mutational effects. However, results based on F1 and F2 estimates were consistent (Fig 3).

We note here that because irradiated individuals are likely to repair some of the radiation-induced DNA damage, and the efficacy of this repair is likely to depend on the condition of the individual (Sharp and Agrawal 2012; Berger et al. 2017), our design only allowed us to test for difference in the temperature-dependence, but not for mean differences, in mutational effects between our two evolution regimes. This is because by raising the F0 generation at 36˚C, control lines will have been more stressed at the time of repair, and thus, could have a greater mutation load because they carried more mutations, and not because these mutations had greater fitness effect when measured on their genetic background (Fig. 3). This hypothesis is supported by control lines having lower mutation loads if raised at the ancestral temperature prior to irradiation (Berger et al. 2017). To estimate the effects of elevated temperature on mutational fitness effects in the two genetic backgrounds, we analysed the number of offspring produced as a Poisson response, using generalized linear mixed effects models, testing for interactions between radiation treatment, assay temperature and evolution regime. We included each individual observation as a random effect to account for over-dispersion in the data. Mutation load is formally quantified as offspring production in irradiated lineages *relative* to corresponding controls. To better illustrate the results we therefore also ran Bayesian analyses using the MCMCglmm package (Hadfield 2010) with the same model structure, but assuming a normally distributed response, and calculated the posterior estimates of mutation load (**Δ𝝎 = 1- 𝝎_IRR_/𝝎_CTRL_** ) directly from these models (Fig. 3). The MCMC resampling ran for 1.000.000 iterations, preceded by 500.000 burn-in iterations that were discarded. Every 1000^th^ iteration was stored, resulting in 1000 independent posterior estimates from each model. We used weak priors for the random effects as recommend in (Hadfield 2010).

**Meta-analysis of selection on de novo mutation in benign and stressful environments**

We looked for studies that had measured fitness effects of de novo mutations in at least two environments, of which one had been labelled stressful relative to the other by the researchers of the study. We started by extracting data from studies reported in two earlier reviews on mutational fitness effects (Martin and Lenormand 2006; Agrawal and Whitlock 2010). We then used Google Scholar to search the literature citing these papers. In addition we also made own searches including the search terms “mutation”, “selection”/”fitness” and “environment”/”stress/”temperature””. We collated selection coefficients along with their standard errors from raw data, tables or figures from the original publications. In all but two cases analysed this labelling was correct in the sense that fitness estimates, based either on survival, reproductive output or population growth rate, were lower in the environment labelled as stressful. In the remaining two cases, the temperature assigned as stressful did not have an effect on the nematode *Caenorhabditis briggsae* (Baer et al. 2006); these estimates were therefore excluded when analysing effects of environmental stress on selection (Fig. 4), but included when analysing the effect of temperature (Fig 5). The studies measured effects of mutations accrued by mutation accumulation, mutagenesis, or targeted insertions/deletions, relative to wild-type controls. We found a few cases that were excluded from analysis since it seemed likely that the protocol used to accrue mutations (mutation accumulation at population sizes >2) may have failed to remove selection, biasing subsequent comparisons of mutational fitness effects across environments. In total we retrieved 100 paired estimates of selection from 28 studies and 11 organisms, spanning unicellular viruses and bacteria to multicellular plants and animals (summary in Supplementary 3). Ultimately, three of these studies (and six paired estimates of selection) were discarded since selection coefficients in both the benign and stressful environment were ≈ 0 and could not be analyzed further.

An estimate controlling for between-study variation was calculated by taking the log-ratio of the cumulative fitness effect of the induced mutations at stressful relative to corresponding benign conditions in each study: LOG_e_[$\boldsymbol{\Delta\omega}$_stress_/**Δ𝝎**_benign_], where $\boldsymbol{\Delta\omega}$ **= 1 – 𝝎^mutant^/𝝎^CTRL^.** Hence, a ratio above (below) 0 indicates stronger (weaker) selection against mutations under stress. We used both REML and Bayesian linear mixed effects models to estimate if log-ratios differed from 0 for three levels of environmental stress: cold temperature, warm temperature, and other types of stress pooled (Table SI 3.1a), as well as for the total effect of stress averaged across all studies. We also tested if log-ratios differed between the three types of abiotic stress. All models included stress-type, mutation induction protocol and fitness estimate as main effects, although effects of the latter two were never significant. We included study organism and study ID as random effects. Additionally, study organism was crossed with stress type to control for species variation and phylogenetic signal. To further explore large scale signals in the data we performed an analysis including a fixed factor encoding uni- or multicellularity, which was crossed with stress type, allowing us to test for differences in selection between the two groups.

Using the 40 estimates that compared the strength of selection across temperatures, we partitioned the effect of i) temperature stress; quantified as the reduction in mean fitness at the stressful temperature relative to the benign temperature (Table S3.1a), and ii) that of temperature itself; quantified as the linear (1^st^ polynomial coefficient) and non-linear (2^nd^ polynomial coefficient) effect of the magnitude and direction of the temperature shift: T_stress_ - T_benign_. We included stress and temperature as the two fixed effect covariates, and study organism and study ID as random effects. Study organisms were also allowed to have random slopes for the temperature effect to control for between-species variation in the temperature dependence. Again we added a fixed effect encoding uni- or multicellularity crossed by the temperature covariate to test if the two groups differed in the temperature dependence of mutational fitness effects.

To weight each estimate’s contribution to the final meta analytic results by its sampling variance, we passed the standard error (SE) of each log-ratio to MCMCglmm using the idh(SE):units command. The standard errors were approximated using laws of error propagation for ratios, but since this technique is known to heavily inflate standard errors when the denominator approaches zero (Fieller 1954), we simulated unidirectional standard errors for the 10 log-ratios for which **Δ𝝎**_benign_ (i.e. the denominator) was smaller than 1.96 SE. This was done by drawing 10.000 samples of $\boldsymbol{\Delta\omega}$_stress_ and **Δ𝝎**_benign_ from a normal distribution defined by their reported mean and standard error and then discarding the 50% of the simulations in which values of **Δ𝝎**_benign_ were below its mean. We then approximated the unidirectional (downwards) error of the log-ratio based on the remaining simulations by calculating the average deviation from the mean log-ratio. Note here that this unidirectional error corresponds directly to whether the log-ratio was significantly different from zero or not (i.e. giving the uncertainty downwards for positive ratios). We present a funnel plot depicting the precision (1/SE) and mean log-ratio in Supplementary 3 (Fig. S3.5).

In all models, the MCMC resampling ran for 2.000.000 iterations, preceded by 500.000 burn-in iterations that were discarded. Every 2000^th^ iteration was stored, resulting in 1000 independent posterior estimates from each model. We used standard priors for the fixed effects and weak priors for the random effects. Variance for the random effect incorporating the within-study standard errors was fixed to 1. All specifications and output from these Bayesian mixed models are presented in Table S1.b.

Friedberg, Errol C., Graham C. Walker, Wolfram Siede, and Richard D. Wood. 2005. *DNA Repair and Mutagenesis*. American Society for Microbiology Press.

Hadfield, Jarrod D. 2010. ‘MCMC Methods for Multi-Response Generalized Linear Mixed Models: The MCMCglmm R Package’. *Journal of Statistical Software* 33 (2): 1–22.

Martin, Guillaume, and Thomas Lenormand. 2006. ‘The Fitness Effect of Mutations across Environments: A Survey in Light of Fitness Landscape Models’. *Evolution* 60 (12): 2413–2427.

Martinossi-Allibert, I., G. Arnqvist, and D. Berger. 2017. ‘Sex-Specific Selection under Environmental Stress in Seed Beetles’. *Journal of Evolutionary Biology* 30 (1): 161–73. https://doi.org/10.1111/jeb.12996.

Rogell, Björn, William Widegren, Lára R. Hallsson, David Berger, Mats Björklund, and Alexei A. Maklakov. 2014. ‘Sex-Dependent Evolution of Life-History Traits Following Adaptation to Climate Warming’. Edited by Charles Fox. *Functional Ecology* 28 (2): 469–78. https://doi.org/10.1111/1365-2435.12179.

Shabalina, Svetlana A., Lev Yu Yampolsky, and Alexey S. Kondrashov. 1997. ‘Rapid Decline of Fitness in Panmictic Populations of Drosophila Melanogaster Maintained under Relaxed Natural Selection’. *Proceedings of the National Academy of Sciences* 94 (24): 13034–39.

Sharp, N. P., and A. F. Agrawal. 2012. ‘Evidence for Elevated Mutation Rates in Low-Quality Genotypes’. *Proceedings of the National Academy of Sciences* 109 (16): 6142–46. https://doi.org/10.1073/pnas.1118918109.
