## Supplementary Information for "Elevated temperature increases genome-wide selection on de novo mutations"

**for:**

**Short title:** Temperature dependent mutational fitness effects

**Key words:** temperature, environment, adaptation, selection, mutation, biodiversity, climate change, enzyme-kinetics, protein stability

**Classification:** Biological Sciences; Evolution

**Supplement 1:** Enzyme-kinetic models of temperature dependent mutational fitness effects

**Supplement 2:** Experimental Evolution of temperature dependent mutational fitness effects

**Supplement 3:** Meta-analysis of temperature dependent mutational fitness effects

**Supplementary Information 1:**

***Biophysical models predict temperature-dependent selection on mutations***

**S1.1: Choice and effect of *Δ***
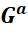
 **and *ΔΔ***
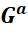
***.***

We explored mutational effects on the activation energy of catalytic rate, *ΔΔ*
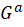
, using data in the SKEMPI 2.0 database, containing Gibbs energies for protein binding of 1844 mutant and corresponding wildtype proteins at standard temperature, typically 25˚C (Jankauskaite et al. 2019). The dataset reflects the motivation and selectivity of researchers of the original studies, interested in understanding protein function and design (Jankauskaite et al. 2019), and hence, do not represent the effect of random mutations on binding. Rather, this data reflect large effect mutations occurring at the binding site itself. Indeed, the mean mutational effect on *ΔΔ*
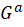
in this data is equal to 0.34 kcal/mol, which, according to Eq. 6 would reduce fitness by more than 40% assuming a 1:1 relationship between enzymatic reaction rate and fitness. Therefore, we first used the data only as a rough guide to approximate a range of plausible values of *ΔΔ*
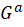
that could parameterize eq.6, used to predict the overall temperature dependence of mutational effects in real organisms and empirical data (see main text). Second, we evaluate how strong the temperature dependence of mutational effects on catalytic rate can maximally be both analytically and based on a smaller subset of mutants measured for both the enthalpic (*ΔΔ*
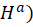
 and temperature-dependent entropic (*ΔΔ*
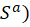
 term of the Gibbs energy for protein binding (see further below).

**Plausible values of *ΔΔ***
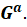
 Given that the binding site itself typically comprises about 10% of the whole protein, the calculated fitness effects in the SKEMPI data could be weighted by this proportion if assuming that mutations outside the binding site would have negligible effects on binding, and that the SKEMPI data do not overestimate effects of mutations occurring within binding sites. If so, the mean effect of a non-synonymous mutation would cause a mean fitness cost of ca. 5% (s ≈ 0.05) according to Eq. 6. This approximation is very rough, and indeed, still seems to generate relatively large (and skewed) fitness effects. We therefore parameterized eq. 6 by taking the pragmatic approach of choosing parameter values of *ΔΔ*
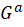
 that generated negative selection coefficients against non-synonymous mutations that were in reasonable agreement with those observed for organisms living at benign temperatures (s ≈ 0-0.01, T = 298°K, see Figure 1), using eq. S1.1b below. We note here that future theoretical and empirical work is needed to better understand and characterize the functional basis for mutational fitness effects on enzyme catalysis (Hochachka & Somero 2002, Echave & Wilke 2017, Serihojos et al. 2017, see Rodriquez et al. 2016 for an example). Combining statistical thermodynamics with knowledge of species ecology and its effects on metabolism would further improve predictions of how mutational effects on enzyme catalysis translates into fitness effects in complex organisms living in their natural environment. For a more general discussion see (Hochachka & Somero 2002, Clarke 2004, Angilletta 2009).

**Temperature dependence via *ΔΔ***
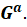
 Here we analytically show that temperature dependence from mutations on activation energy is expected to be very weak and negative. Based on the standard formulation of enzyme kinetics (Eqs. 1 & 4, main text), selection on mutations affecting the activation energy at a given temperature equals:

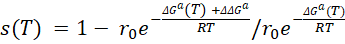
,	(Eq. S1.1a)

Where, R is the universal gas constant = 0.002 kcal/mol, and T is temperature measured in degrees Kelvin. Solving for the mutational effect yields:

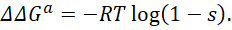
(Eq. S1.1b)

This standard formulation of enzyme catalytic rate makes two predictions regarding mutational fitness effects through changes in the activation energy, *ΔΔ*
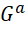
. First, the selection coefficient is independent of the original value of *Δ*
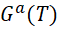
, and therefore also insensitive to temperature-induced changes in *ΔG^a^* (Fig. S1.1A). Second, rearranging and exponentiating eq. S1.1b yields:

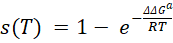
(Eq. S1.1c)

Which, on the basis that: *∆∆G* = ∆∆H + *TΔΔ*S, can be re-written as:

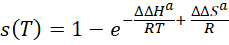
(Eq. S1.1d)

Eqs. S1.1c and S1.1d show that, contrary to the net increase in mutational fitness effects at elevated temperature observed in data (main text), fitness effects of mutations affecting the activation energy, *ΔΔ*
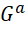
*,* should show a negligible *negative* temperature-dependence for any ecologically relevant temperature change (i.e. 273-313K) (Fig. S1.1A).

**Exploring effects of mutation on *ΔΔ***
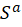
 **in the SKEMPI data.** We evaluated how large mutational effects on the entropy of the Gibbs energy (i.e. *ΔΔ*
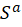
) are relative to enthalpy (*ΔΔ*
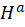
) changes for 443 available mutants in the SKEMPI data. Simultaneous estimation of entropic and enthalpic terms are associated with statistical interdependencies and considerable measurement error (Cornish-Bowden 2002, Chodera & Mobley 2013, Khrapunov 2018), and the SKEMPI data is unfiltered with regards to such problems (Jankauskaite et al. 2019). Nevertheless, the data can be used to generate predictions of mean enthalpic and entropic mutational changes by averaging across entries (Fig. S1.1B). To reduce the inherent bias in the original data, we first removed entries where temperature-independent enthalpy changes (*ΔΔ*
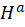
) between wildtype and mutant exceeded 10 kcal/mol, potentially reflecting large measurement errors or mutations with extraordinary strong effects. Additionally, we removed 45 entries where the wildtype and mutant had been measured at different experimental temperatures. This resulted in 366 entries left for analysis.

The mean mutational effect on *ΔΔ*
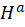
 equals 1.03 (sd = 3.5) kcal/mol, whereas the effect on *ΔΔ*
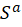
is highly variable, not significantly different from 0, and equal to -0.00016 (sd=0.01). We note that the variability in these estimates will be greatly overestimated due to correlated measurement errors (Chodera & Mobley 2013). This translates to a mean contribution of the entropy term equal to -0.049 kcal/mol at standard temperature (T = 298˚K). Hence, although both these terms contribute to the overall mutational effect on the Gibbs energy by: *ΔΔ*
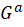
 = *ΔΔ*
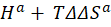
, the destabilizing contribution from the enthalpic term is greater on average. While mutational effects on the activation energy are not predicted to generate strong temperature dependence on average (Fig. S1A, B), specific mutations in the SKEMPI data could, which would tend to cause variability across experimental studies. Importantly, the temperature dependence generated from mutations affecting the activation energy is predicted to be weakly negative on average, the opposite pattern to that predicted for mutations affecting folding and what is observed in data (Figs. 4 & 5, main text).

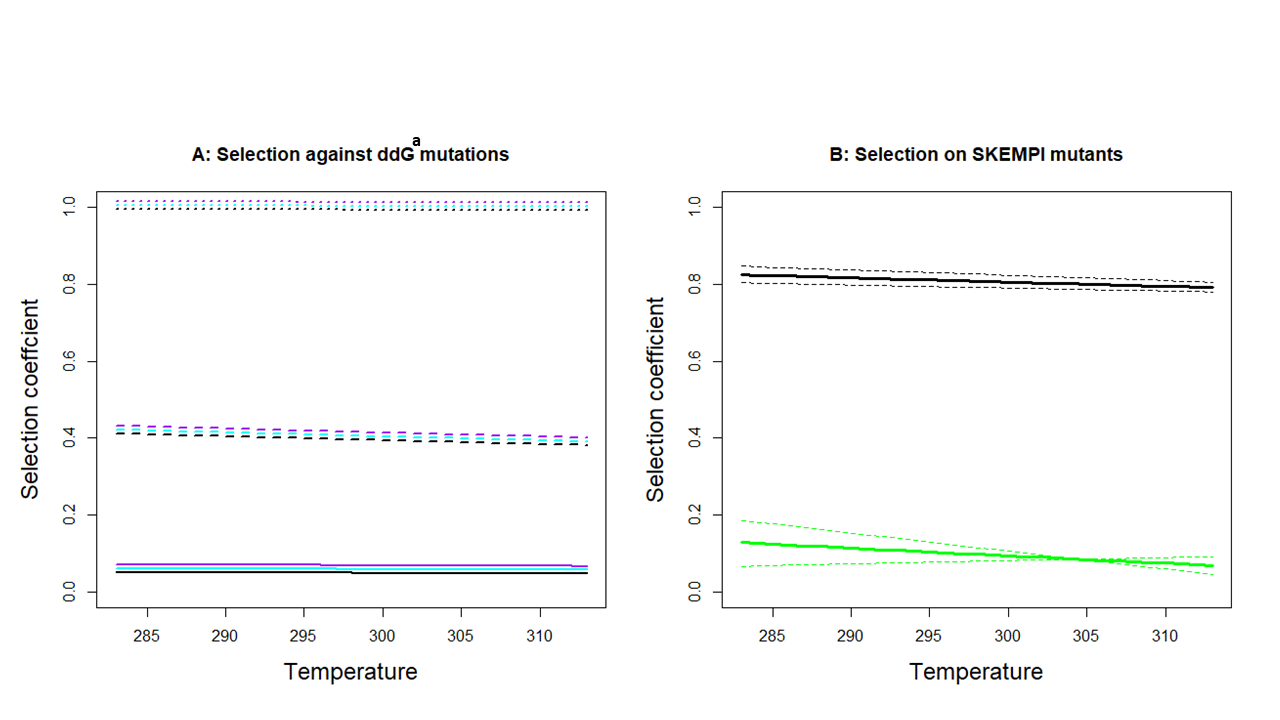

**Figure S1.1: Predicted temperature-dependence of selection against mutations affecting catalytic rate.** In **A**) selection against three hypothetical mutants with moderate (full lines;
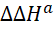
 = 0.03), strong (hatched lines;
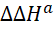
 = 0.3) or very strong (dotted lines;
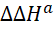
 = 3) effects. Temperature dependence is negligible and independent of the original activation energy of the wildtype (black: *Δ*
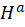
 = 20, *Δ*
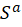
 = 0.001; blue: *Δ*
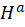
 = 20, *Δ*
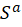
 = 0.1; purple: *Δ*
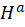
 = 10, *Δ*
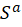
 = 0.001). Colored lines have been shifted horizontally for illustration. In **B**) predicted mean selection on SKEMPI mutants scored for both *ΔΔ*
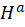
 and *ΔΔ*
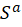
 (mean = full lines, 95% CI = hatched lines). Black lines depict predictions based on all 366 mutants and green lines for the subset of 90 mutants with the weakest effect on binding.

Angilletta, Michael J. *Thermal Adaptation: A Theoretical and Empirical Synthesis*. Oxford Biology. Oxford ; New York: Oxford University Press, 2009.

Bershtein, Shimon, Adrian WR Serohijos, and Eugene I Shakhnovich. 2017. “Bridging the Physical Scales in Evolutionary Biology: From Protein Sequence Space to Fitness of Organisms and Populations.” *Current Opinion in Structural Biology*, Folding and binding • Proteins: Bridging theory and experiment, 42 (February): 31–40. <https://doi.org/10.1016/j.sbi.2016.10.013>.

Chodera, John D., och David L. Mobley. ”Entropy-Enthalpy Compensation: Role and Ramifications in Biomolecular Ligand Recognition and Design”. *Annual Review of Biophysics* 42, nr 1 (2013): 121–42. <https://doi.org/10.1146/annurev-biophys-083012-130318>.

Clarke, A. 2004. “Is There a Universal Temperature Dependence of Metabolism?” *Functional Ecology* 18 (2): 252–56. <https://doi.org/10.1111/j.0269-8463.2004.00842.x>.

Cornish-Bowden, Athel. ”Enthalpy—Entropy Compensation: A Phantom Phenomenon”. *Journal of Biosciences* 27, nr 2 (01 mars 2002): 121–26. <https://doi.org/10.1007/BF02703768>.

Hochachka, Peter W., and George N. Somero. 2002. *Biochemical Adaptation: Mechanism and Process in Physiological Evolution*. Oxford University Press.

Jankauskaitė, J., Jiménez-García, B., Dapkūnas, J., Fernández-Recio, J. & Moal, I. H. SKEMPI 2.0: an updated benchmark of changes in protein–protein binding energy, kinetics and thermodynamics upon mutation. *Bioinformatics* **35**, 462–469 (2019).

Khrapunov, Sergei. ”The Enthalpy-entropy Compensation Phenomenon. Limitations for the Use of Some Basic Thermodynamic Equations”. *Current protein & peptide science* 19, nr 11 (2018): 1088–91. <https://doi.org/10.2174/1389203719666180521092615>.

Rodrigues, João V., Shimon Bershtein, Anna Li, Elena R. Lozovsky, Daniel L. Hartl, and Eugene I. Shakhnovich. 2016. “Biophysical Principles Predict Fitness Landscapes of Drug Resistance.” *Proceedings of the National Academy of Sciences* 113 (11): E1470–78. <https://doi.org/10.1073/pnas.1601441113>.

**S1.2: Temperature-dependent selection on mutations affecting only enthalpy, only entropy, or both.**

To explore if our qualitative results were dependent on whether mutational effects on folding (*∆∆G^f^*) were induced via the enthalpic (*∆∆H^f^*) or entropic term (*∆∆S^f^*), we estimated the temperature dependence of the mean mutational fitness effect (i.e. the mean selection coefficient) and mutational variance (i.e. the squared selection coefficient) in three simulated scenarios: 1) effects only via *∆∆H^f^* (as in Fig. 1 of the main text), 2) effects only via *∆∆S^f^*, and 3) effects via both *∆∆H^f^* and *∆∆S^f^* with entropy-enthalpy compensation. In all scenarios, effects on folding were explored in a single protein with a stability, *∆G^f^* = -6 kcal/mol K^-1^ and *∆S^f^* = 0.25 or 1.0 kcal/mol K^-1^ based on values for typical proteins (Hochachka & Somero 2002, Dill et al. 2011). For simplicity we here assumed mutational effects on activation energy to be zero (*∆∆G^a^* = 0).

In scenarios 1 and 2 we first drew mutational effects on folding at 298K from the empirical normal distribution (*∆∆G^f^* mean = 0.9, sd = 1.7 kcal/mol, see main text) and then let this effect be fully accounted for by either the enthalpic or entropic term at 298K (*∆∆G^f^* ≈ *∆∆H^f^*+ *T∆∆S^f^* for K_283-313_). For scenario 3 we again drew mutational effects on stability from the empirical distribution of *∆∆G^f^*. We then set the entropic (scenario 3a) or enthalpic (scenario 3b) effect to be 5 times greater on stability at 298K, resulting in 5 times greater mean effect and 25 times more variance in the simulated effect. The extra change in folding energy generated by the excess variance in the entropic (enthalpic) term was then compensated by a corresponding (antagonistic) change in the enthalpic (entropic) term so that the resulting *∆∆G^f^* at 298K still corresponded to the observed empirical distribution.

In scenario 3 we assumed that enthalpic and entropic terms could vary substantially more than mutational effects on the Gibbs energy, more so than what is typically seen in data. We also note here that the extent of enthalpy-entropy compensation is highly disputed and most empirical estimates are flawed by correlated measurement errors (reviewed in: Cornish-Bowden 2002, Chodera & Mobley 2013, Khrapunov 2017). However, our aim here was to produce an extreme scenario to evaluate whether mutations affecting the temperature dependence of the Gibbs energy itself could be large enough to offset the increase in mutational fitness effects predicted at elevated temperature. This could happen if mutations confer reduced temperature sensitivity on average (by reducing the term *T∆∆S^f^*), which is the case in scenario 3b (bottom row in Fig. S1.2). However, as our simulations show, this effect is weak and cannot override the general temperature induced increase in the entropic energy of the original protein configuration. That is, the change in *∆S^f^* in the mutant that can be generated while being (reasonably) consistent with observed values of *∆∆G^f^* is small relative to the entropic term in the wildtype, as can be expected from single point mutations in a protein containing hundreds of amino acids. As laid out in the main text, the general effect of temperature to increase entropy (i.e. *T∆S^f^* is always positive for both wildtype and mutant) is predicted to cause marginally stable proteins to rapidly unfold as temperature increases, and this general temperature dependence is in turn predicted to cause mutational fitness effects to increase at elevated temperature (Fig. S1.2).

Chodera, John D., och David L. Mobley. ”Entropy-Enthalpy Compensation: Role and Ramifications in Biomolecular Ligand Recognition and Design”. *Annual Review of Biophysics* 42, nr 1 (2013): 121–42. <https://doi.org/10.1146/annurev-biophys-083012-130318>.

Cornish-Bowden, Athel. ”Enthalpy—Entropy Compensation: A Phantom Phenomenon”. *Journal of Biosciences* 27, nr 2 (01 mars 2002): 121–26. <https://doi.org/10.1007/BF02703768>.

Dill, Ken A., Kingshuk Ghosh, och Jeremy D. Schmit. ”Physical Limits of Cells and Proteomes”. *Proceedings of the National Academy of Sciences* 108, nr 44 (01 november 2011): 17876–82. <https://doi.org/10.1073/pnas.1114477108>.

Hochachka, Peter W., och George N. Somero. *Biochemical Adaptation: Mechanism and Process in Physiological Evolution*. Oxford University Press, 2002.

Khrapunov, Sergei. ”The Enthalpy-entropy Compensation Phenomenon. Limitations for the Use of Some Basic Thermodynamic Equations”. *Current protein & peptide science* 19, nr 11 (2018): 1088–91. <https://doi.org/10.2174/1389203719666180521092615>.

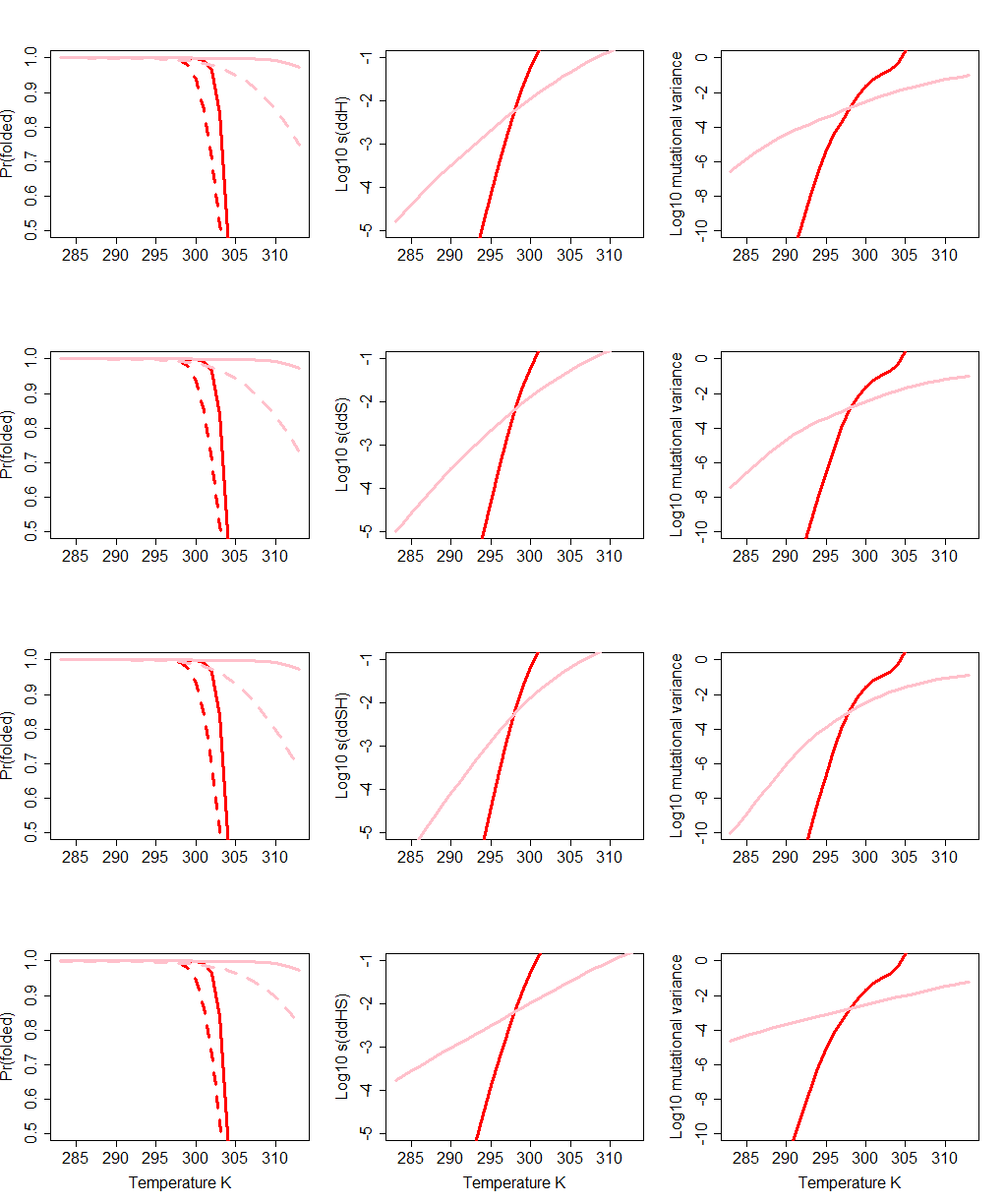

**Figure S1.2: Effects of mutational effects on protein stability via entropic and enthalpic contributions.** Effects on (left to right) the proprtion of folded protein, the mean strenght of purifying selection, and mutational variance in fitness for four different scenarions (top to bottom): Scenario 1 – effects on enthalpy only (as used in the examples in the main text), Scenario 2 - effects via entropy only, Scenario 3a – enthalpy-entropy compensation with entropic effects being destabilizing on average at 298K, and Scenario 3b – enthalpy-entropy compensation with enthalpic effects being destabilizing on average at 298K. In the left panels the wildtype protein is represented by full lines and the mutant protein by dashed lines. Red lines: *∆S^f^* = 1.0 kcal/mol K^-1^, pink lines: *∆S^f^* = 0.25 kcal/mol K^-1^.

**S1.3: The distribution of mutational fitness effects as a function of temperature and protein stability**

**Fig. S1.3.** Distribution of selection coefficients on folding mutations, *ΔΔ*
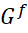
, in three single proteins with different stabilities, *Δ*
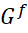
, (unstable = -6, moderate = -9, robust = -12) at three temperatures. Most mutations are deleterious, and the overall strength of selection is strongest for the least stable protein and increases with temperature for all three proteins. The fraction of highly deleterious as well as positively selected mutations increase at warm temperature. Selection coefficients were generated using equation (6) and sampling from the empirical normal distribution of *ΔΔ*
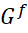
 (see main text).

20 ^o^C

30 ^o^C

40 ^o^C

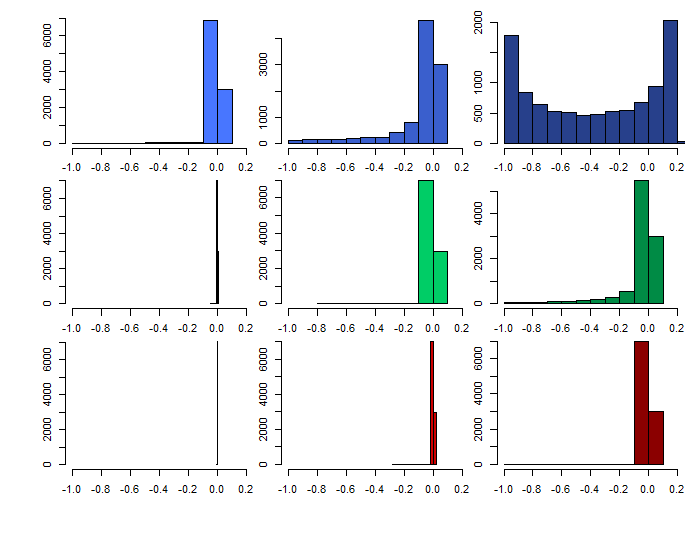

*ΔG* = -6

*ΔG* = -9

Selection coefficent

*ΔG* = -12

**S1.4: Effects of thermal sensitivity of entropy terms for folding, *Δ***
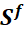
***.***

Differences in the temperature dependence of the entropy term of the Gibbs energy of folding, *ΔS^f^,* among species could potentially explain some of the variance in the temperature dependence of mutational effects seen among studies in our meta-analysis (see also Supplementary S1.2). Here we explored this effect across whole protein ensembles by sampling of the empirical distributions of *ΔG^f^* and *ΔΔG^f^* (see main text). Indeed, this predicted effect is seen in the effects of folding mutations, particularly evident at low temperatures when we consider the simultaneous effects of *ΔΔG^f^* and *ΔΔG^a^* mutations. Nonetheless, our key prediction that the strength of selection against mutations increases with temperature is robust to model parameterization (Fig. S1.4).

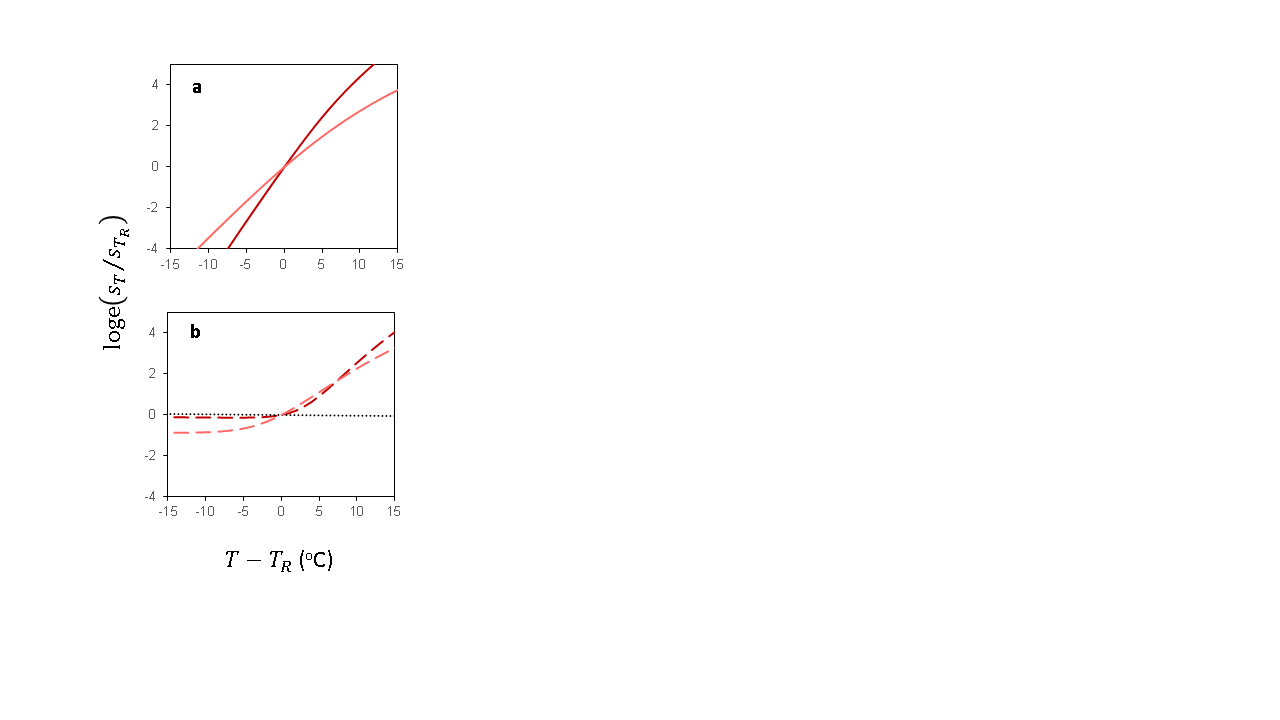

**Fig S1.4:** Relative strength of selection against a mutation as a function of temperature, plotted relative to a (benign) standardized reference temperature defined as:
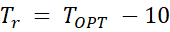
 ^o^C (as in fig 1d). Here we compare the response of a temperature sensitive genotype (*ΔS^f^* = 0.3; dark red line) to a more insensitive genotype (*ΔS^f^* = 0.2; pink line). A higher *ΔS^f^* leads to greater sensitivity to a given temperature change. For clarity results are only given for a ’warm-adapted’ species (*ΔG^f^_K=298_* = -12) as the results are the same for other ecotypes (as in fig 1c). In a), *ΔΔG^a^* = 0; in b), *ΔΔG^a^* yields a 10^-3^ loss in relative fitness at
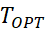
 (long-dash lines). In b) the black dotted line represents the response of a mutant affecting only *ΔG^a^.*

**S1.5. An alternative model of temperature-dependent mutational fitness effects based on metabolic flux and protein toxicity**

The effect of protein stability on metabolic flux through a linear metabolic pathway can be written as:

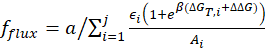
,	(Eq. S1.4a)

where
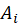
 is the abundance of protein
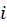
,
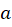
 is a normalisation constant equal to the abundance of the input metabolite across
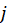
genes (i.e.
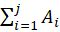
) and
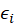
 is the enzyme activity of protein
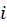
 (Echave & Wilke 2017). A review of protein abundances in cells suggests the total abundance of cell proteins typically ranges from 10^6^ to 10^9^ copies (Milo 2013). Models of
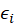
 for actual proteins are still lacking (Echave & Wilke 2017), so for simplicity we set this value to 1 for all
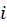
. We note that mutational effects on enzyme activation also could be modelled but this would not generate any strong temperature dependence (see main text and Supplementary S1.1). Note also that the temperature-dependence of protein misfolding here arises from the terms
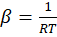
 and
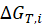
 =
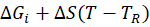
 (see main text for further details).

Most biophysical models of protein evolution directly map the probability protein folding to fitness. However, fitness might to large extent be determined by cytotoxicity associated with the amount of misfolded protein in a cell (Drummond & Wilke 2008), which can be modelled as (Echave & Wilke 2017):

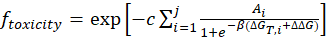
,	(Eq. S1.4b)

where
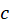
 is a measure of toxicity per misfolded protein and the sum runs over all genes
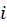
 in the organism's genome and has been estimated to 10^-4^ (Drummond & Wilke 2008, Serohijos & Shakhnovich 2013). Fitness is calculated here as the product of
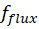
 and
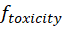
.

This alternative model corroborates our main findings by reproducing qualitatively and quantitatively similar responses as the biophysical model presented in the main text. While this flux-toxicity model is intrinsically more difficult to parameterize, it provides additional mechanistic insights that potentially can explain variation in the predicted universal increase in mutational fitness effects with temperature through the level of protein expression of the mutated gene. In figure S1.5 we illustrate the consequences of a mutation (
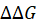
 = *∆∆H* = 0.9) in a gene with a given
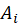
 within a given metabolic pathway consisting of 10 genes.

Drummond, D. Allan, & Claus O. Wilke. (2008). Mistranslation-Induced Protein Misfolding as a Dominant Constraint on Coding-Sequence Evolution. *Cell* 134: 341–52.

Echave, J., & Wilke, C. O. (2017). Biophysical Models of Protein Evolution: Understanding the Patterns of Evolutionary Sequence Divergence. *Annual review of biophysics*, *46*, 85-103.

Milo R. (2013). What is the total number of protein molecules per cell volume? A call to rethink some published values. *BioEssays : news and reviews in molecular, cellular and developmental biology*, *35*(12), 1050-5.

Serohijos, A. W., & Shakhnovich, E. I. (2013). Contribution of selection for protein folding stability in shaping the patterns of polymorphisms in coding regions. *Molecular biology and evolution*, *31*(1), 165-76.

**Fig S1.5. Relative strength of selection against a mutation as a function of temperature predicted by an alternative biophysical model of protein folding based on flux and cytotoxicity in a linear metabolic pathway.**

Upper panels depict the selection coefficients, the lower panels the log ratio of selection coefficients with temperature. The temperature dependence of mutational effect is always positive. However, the strength of this relationship is subject to the gene’s protein expression level
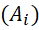
. Genes giving rise to more abundant proteins are subject to greater selection pressure via deleterious effects of cytotoxicity. Results shown for
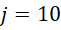
 genes, with
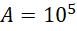
 proteins for the 9 genes with the wildtype allele and A =
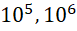
or
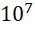
 for the gene carrying the mutation (light grey to black lines). The 10 genes encode proteins with stabilties ranging stepwise from -5 to -14 and shown are results for a mutation in a gene encoding a less stable (*Δ*G = -6, left panels) and more stable protein (*Δ*G = -7, right panels). As for the biophysical model presented in the main text, mutational fitness effects are stronger in less stable proteins.

Less stable protein (
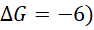

More stable protein (
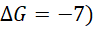

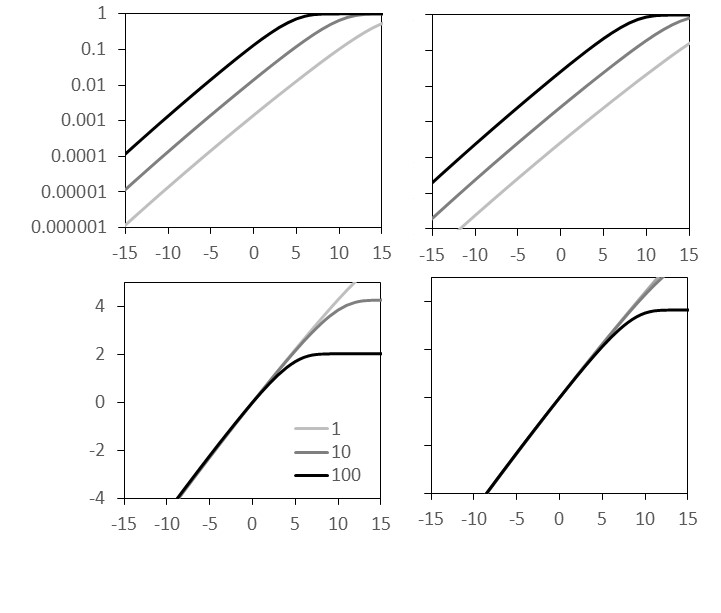

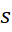

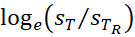

10^5^

10^6^

10^7^

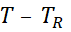
(^o^C)

**Supplementary Information 2**

***Experimental evolution shows evidence for temperature-dependent selection on induced mutations in seed beetles***

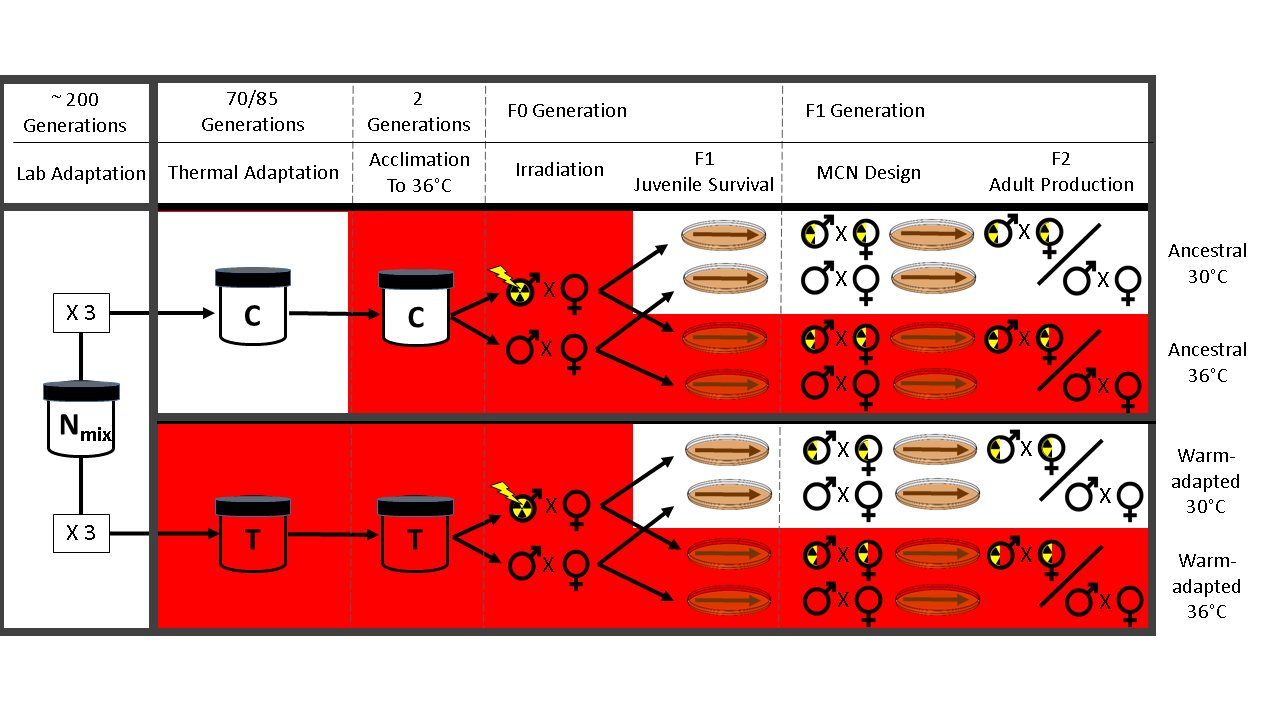

**Fig SI 2.1: Broad schematic overview of the experimental design used to induce and measure mutations in Callosobruchus maculatus (see *Methods* for details).**

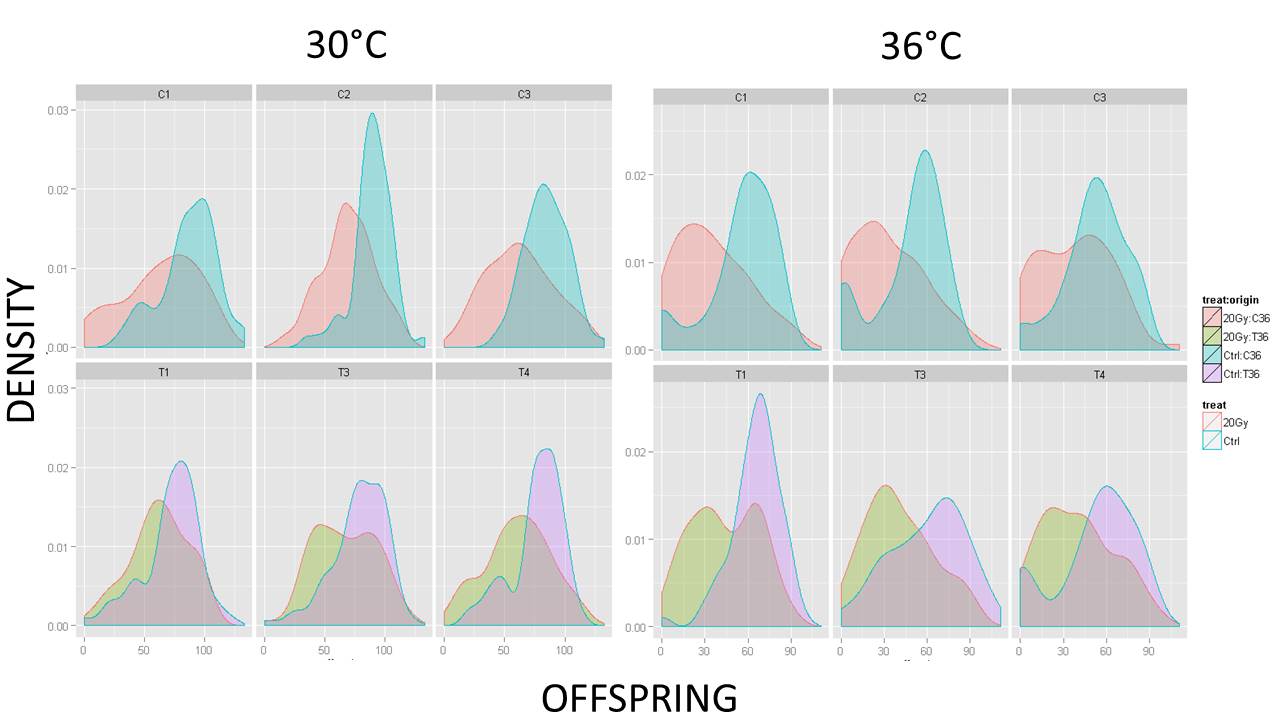

**Fig SI 2.2:** **Density distribution of lifetime F2 offspring production in the irradiated and control lineages for each line and temperature** (C = ancestral lines, T = warm-adapted lines).

**Supplementary Information 3:**

***Meta-Analysis of Mutational Effects***

**SI Table 3.1a:** **Summary of the 100 estimates comparing selection on de novo mutation in benign and stressful environments, from 28 published studies on 11 species**.

Andrew et al. 2015, compared the top and bottom 5 MA lines in terms of fitness across environments to assess mutational effects. However, since ranking was done in the benign environment, the cross-environment comparison is potentially biased (Halligan & Keightley 2009). Here we therefore averaged the 10 MA lines and compared them to available proper controls to estimate selection. For Dandage et al. (2018) selection was calculated from estimates of presence/absence (implying survival) after saturated growth, rather than reported measures of growth rate that discarded absent mutants. In Szafraniec et al. (2001) one measure of the ratio log(Δ𝝎_stress_/Δ𝝎_benign_) was extremely large because selection at the benign temperature was ~0. Therefore, this ratio was set equal to the paired estimate from the same study (=4.31). This is thus a conservative point estimate of the increase in selection at hot temperature. The three studies (6 estimates) from *Arabidopsis thaliana* could not be analyzed further since all selection against mutants in all environments was ~0. Stress in terms of fitness in the stressful relative to the benign environment (𝝎_s_/𝝎_b_) was calculated from the studies except in two cases when it was derived from other studies (these are denoted ^a^ and ^b^ respectively).

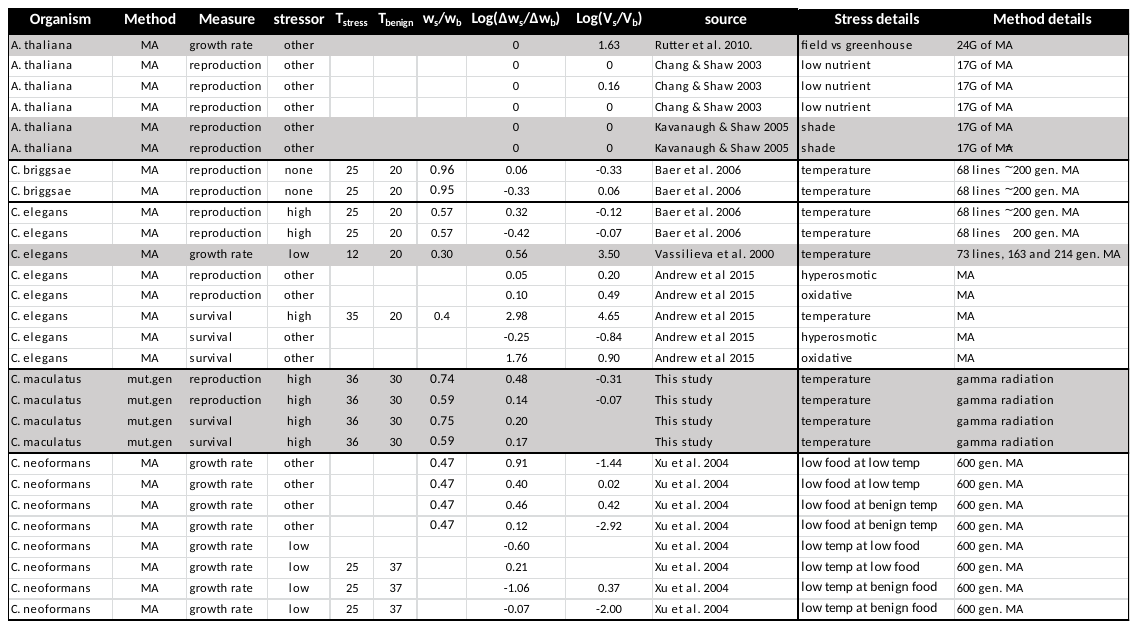

**SI Table 3.1a (Continued)**

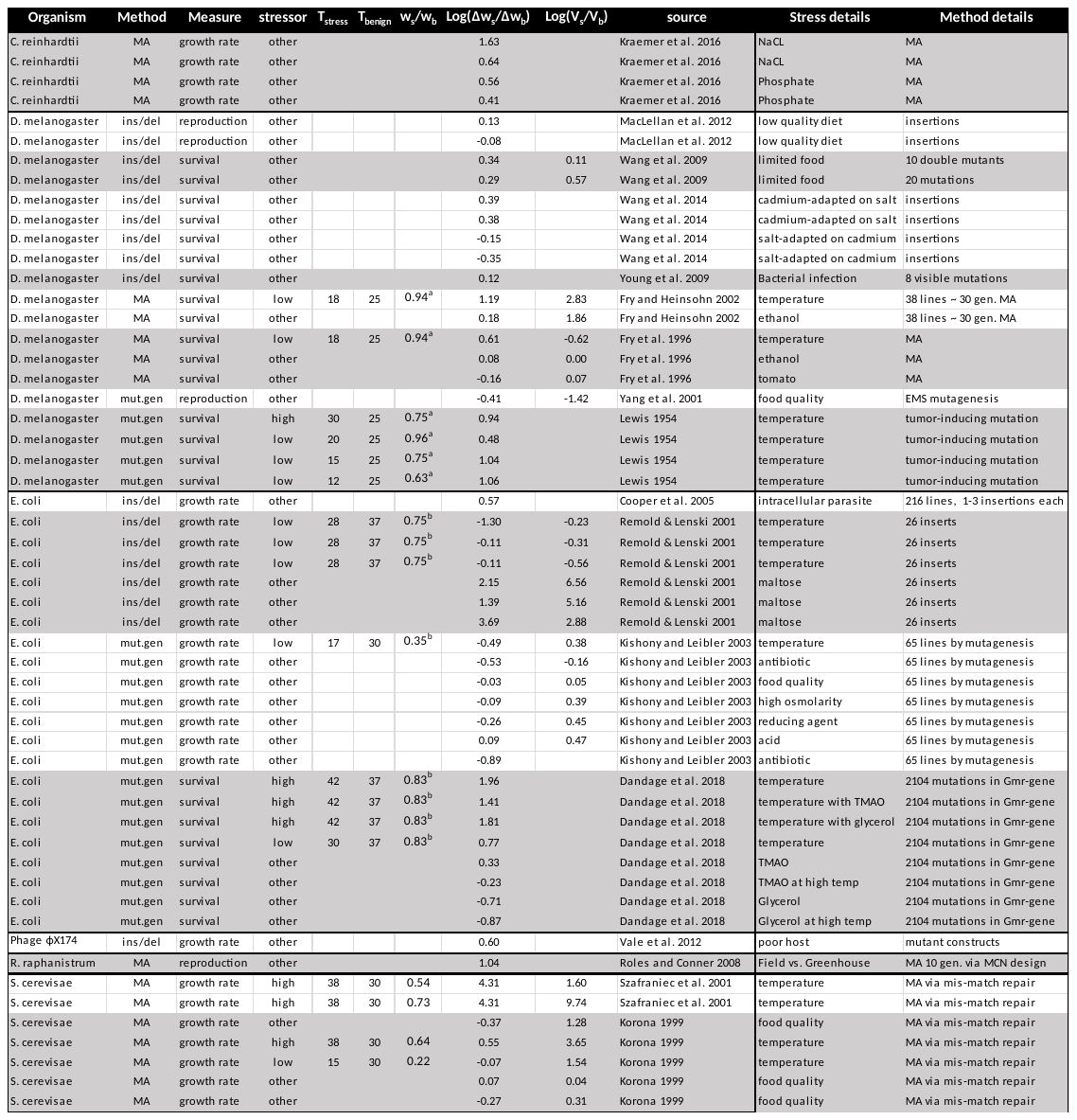

**SI Table 3.1a (Continued)**

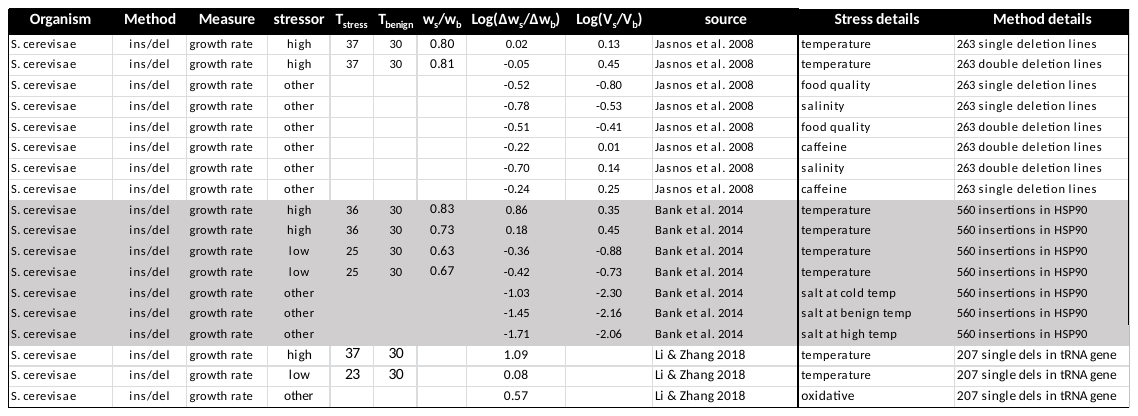

***MA* refers to mutation accumulation, *mut.gen* to mutagenesis (inducing mutations through radioactivity, UV-radiation or chemical mutagens), and *ins/del* refers to insertions or deletions. Fitness measures were provided through estimates of reproduction (e.g. female egg/offspring production), survival (e.g. egg-to-adult survival) or growth rate (propagation through clonal growth; Malthusian fitness). Estimates of relative fitness at benign and stressful temperature for D. melanogaster and E.coli were derived from Schou et al. 2017(^a^) and Bronikowski et al. 2001 (^b^) respectively. For further information, see main text.**

**SI Table 3.1b: MCMCglmm model specifications and output.**

*Note that model results were insensitive to (reasonable) changes in prior variances (V) and strength of belief (nu). Shown are results from the reported models using the weakest priors.*

```
#Effect of different stressors:

prior1 = list(R = list(V = diag(3), nu = 10^-6),
G = list(G1 =  list(V = 1, nu = 10^-6), G2 =  list(V = 1, nu = 10^-6),G3 =  list(V = 1, nu = 10^-6),G4 =  list(V = diag(1), fix = 1)))

metaMC <- MCMCglmm(logratio ~ stress + method + measure ,
random = ~  source + organism + stress:organism + idh(simSE):units,
rcov = ~ idh(stress):units, data = meta,
family = "gaussian", prior = prior1,  nitt=2500000,
slice=TRUE, burnin=500000, thin=2000, verbose = FALSE, pr=F)


DIC: 112.2702

 G-structure:  ~source

       post.mean l-95% CI u-95% CI eff.samp
source    0.3124  0.03372   0.6925     1000

               ~organism

         post.mean  l-95% CI u-95% CI eff.samp
organism    0.0351 1.546e-07   0.2171    1000

               ~stress:organism

                post.mean  l-95% CI u-95% CI eff.samp
stress:organism    0.2757 1.556e-06   0.6304     1243

               ~idh(simSE):units

            post.mean l-95% CI u-95% CI eff.samp
simSE.units         1        1        1        0

 R-structure:  ~idh(stress):units

            post.mean l-95% CI u-95% CI eff.samp
high.units    0.13222  0.01102   0.3910     1000
low.units     0.43468  0.08882   0.9523     1204
other.units   0.09414  0.02883   0.1769     1097

 Location effects: logratio ~ stress + method + measure

                    post.mean l-95% CI u-95% CI eff.samp pMCMC
Intercept(highT)      1.26850  0.56274  2.06746     1000 0.002 **
stresslow            -1.06989 -1.83786 -0.31082     1013 0.004 **
stressother          -1.10143 -1.86363 -0.47493     1000 0.002 **
methodMA              0.14735 -0.49461  0.78263     1000 0.670
methodmutagenesis    -0.48824 -1.41082  0.34263     1000 0.270
measurereproduction  -0.17999 -0.94622  0.54283     1000 0.638
```

measuresurvival	-0.02485 -0.88969 0.64575	1000 1.000

**SI Table 3.1b (continued)**

```
#Effect of different stressors for unicellular and multicellular organisms:

prior5b = list(R = list(V = diag(3), nu = 10^-6),
G = list(G1 =  list(V = 1, nu = 10^-6), G2 =  list(V = 1, nu = 10^-6),G3 =  list(V = 1, nu = 10^-6),G4 =  list(V = diag(1), fix = 1)))

metaMC2 <- MCMCglmm(logratio ~ stress*clade2 + method + measure ,
random = ~  source + organism + stress:organism + idh(simSE):units,
rcov = ~ idh(stress):units, data = metaOK,
family = "gaussian", prior = prior5b,  nitt=2500000,
slice=TRUE, burnin=500000, thin=2000, verbose = FALSE, pr=F)

 DIC: 115.0862

 G-structure:  ~source

       post.mean  l-95% CI u-95% CI eff.samp
source    0.3446 4.582e-07   0.7424     1000

               ~organism

         post.mean  l-95% CI u-95% CI eff.samp
organism   0.04305 1.553e-07    0.233     1000

               ~stress:organism

                post.mean  l-95% CI u-95% CI eff.samp
stress:organism    0.2134 2.236e-07    0.586    1000

               ~idh(simSE):units

            post.mean l-95% CI u-95% CI eff.samp
simSE.units         1        1        1        0

 R-structure:  ~idh(stress):units

            post.mean l-95% CI u-95% CI eff.samp
high.units    0.15749 0.007291   0.4978     1000
low.units     0.44969 0.099279   0.9531     1000
other.units   0.09588 0.034399   0.1808     1000

 Location effects: logratio ~ stress * clade2 + method + measure

                                post.mean l-95% CI u-95% CI eff.samp  pMCMC
Intercept(highT)                   1.5087   0.7696   2.3003   1000.0 <0.001
stresslow                         -1.5743  -2.5227  -0.7178   1000.0  0.002
stressother                       -1.3128  -2.1321  -0.6002   1000.0  0.004
clade2multicellular               -0.6737  -1.9993   0.7693   1000.0  0.310
methodMA                           0.1458  -0.4810   0.8753   1000.0  0.674
methodmutagenesis                 -0.4775  -1.3705   0.4479   1000.0  0.316
measurereproduction               -0.0484  -0.9860   1.0921   1000.0  0.912
measuresurvival                    0.1031  -0.9382   1.0864   1000.0  0.834
stresslow:clade2multicellular      1.4929  -0.2071   2.8327   1000.0  0.058
stressother:clade2multicellular    0.4809  -0.8028   1.9081   1000.0  0.470
```

**SI Table 3.1b (continued)**

```
#Effect of temperature shift versus drop in relative fitness:

prior6 = list(R = list(V = 1, nu = 10^-6),
G = list(G1 =  list(V = 1, nu = 10^-6), G2 =  list(V = 1, nu = 10^-6),G3 =  list(V = 1, nu = 10^-6),G4 =  list(V = diag(1), fix = 1)))


metaMC <- MCMCglmm(logratio ~ poly(deltaT,2) +  rel.fit + measure + method,
random = ~  source + organism + deltaT:organism + idh(simSE):units,
rcov = ~ units, data = tempdata,
family = "gaussian", prior = prior6,  nitt=2500000,
slice=TRUE, burnin=500000, thin=2000, verbose = FALSE, pr=F)


 DIC: 33.08964

 G-structure:  ~source

       post.mean  l-95% CI u-95% CI eff.samp
source    0.8113 9.644e-07    1.959     1309

               ~organism

         post.mean  l-95% CI u-95% CI eff.samp
organism   0.04354 2.226e-07    0.211     1000

               ~deltaT:organism

                post.mean  l-95% CI u-95% CI eff.samp
deltaT:organism   0.00909 2.139e-07  0.04763    1000

               ~idh(simSE):units

            post.mean l-95% CI u-95% CI eff.samp
simSE.units         1        1        1        0

 R-structure:  ~units

      post.mean l-95% CI u-95% CI eff.samp
units   0.08789  0.01648   0.1827     1000

 Location effects: logratio ~ poly(deltaT, 2) + rel.fit + measure + method

                    post.mean l-95% CI u-95% CI eff.samp pMCMC
Intercept(highT)      0.23298 -1.10460  1.60210     1000 0.724
poly(deltaT, 2)1      2.43939  0.98817  3.68054     1000 0.004 **
poly(deltaT, 2)2      2.11348  0.40775  3.76916     1000 0.034 *
rel.fit              -0.09471 -1.56056  1.40884     1000 0.846
measurereproduction   0.26445 -1.11334  1.65191     1115 0.680
measuresurvival       0.38517 -0.96842  1.75283     1000 0.550
methodMA              0.34550 -1.06106  1.64719     1000 0.594
methodmutagenesis     0.16518 -1.63963  1.65221     1000 0.854
```

**SI Table 3.1b (continued)**

```
#removing non-significant terms (relative fitness, method and measure)

metaMC <- MCMCglmm(logratio ~ poly(deltaT,2) ,
random = ~  source + organism + deltaT:organism + idh(simSE):units,
rcov = ~ units, data = tempdata,
family = "gaussian", prior = prior6,  nitt=2500000,
slice=TRUE, burnin=500000, thin=2000, verbose = FALSE, pr=F)

 DIC: 26.66964

 G-structure:  ~source

       post.mean l-95% CI u-95% CI eff.samp
source    0.6432  0.06927    1.391     1293

               ~organism

         post.mean  l-95% CI u-95% CI eff.samp
organism   0.04379 1.502e-07   0.1962     1000

               ~deltaT:organism

                post.mean  l-95% CI u-95% CI eff.samp
deltaT:organism   0.01397 1.756e-07  0.07771     1000

               ~idh(simSE):units

            post.mean l-95% CI u-95% CI eff.samp
simSE.units         1        1        1        0

 R-structure:  ~units

      post.mean l-95% CI u-95% CI eff.samp
units   0.07546  0.01961   0.1556     1000

 Location effects: logratio ~ poly(deltaT, 2)

                 post.mean l-95% CI u-95% CI eff.samp  pMCMC
(Intercept)         0.5765   0.1082   1.0419   1000.0   0.01 *
poly(deltaT, 2)1    2.4173   1.2208   3.5867   1000.0 <0.001 ***
poly(deltaT, 2)2    2.1158   0.5228   3.4328   1108.8   0.01 *
```

**SI Table 3.1b (continued)**

```
#temperature dependence for unicellular and multicellular species

prior6 = list(R = list(V = 1, nu = 10^-6),
G = list(G1 =  list(V = 1, nu = 10^-6), G2 =  list(V = 1, nu = 10^-6),G3 =  list(V = 1, nu = 10^-6),G4 =  list(V = diag(1), fix = 1)))


metaMC <- MCMCglmm(logratio ~ poly(deltaT,2)*clade2  ,
random = ~  source + organism + deltaT:organism + idh(simSE):units,
rcov = ~ units, data = tempdata,
family = "gaussian", prior = prior6,  nitt=2500000,
slice=TRUE, burnin=500000, thin=2000, verbose = FALSE, pr=F)

DIC: 21.64652

 G-structure:  ~source

       post.mean l-95% CI u-95% CI eff.samp
source    0.4214  0.06125    1.062     1000

               ~organism

         post.mean l-95% CI u-95% CI eff.samp
organism   0.06678 1.04e-07    0.172     1000

               ~deltaT:organism

                post.mean  l-95% CI u-95% CI eff.samp
deltaT:organism  0.008231 1.758e-07  0.03797    1000

               ~idh(simSE):units

            post.mean l-95% CI u-95% CI eff.samp
simSE.units         1        1        1        0

 R-structure:  ~units

      post.mean l-95% CI u-95% CI eff.samp
units   0.06796  0.01323   0.1536     1000

 Location effects: logratio ~ poly(deltaT, 2) * clade2

                               post.mean l-95% CI u-95% CI eff.samp  pMCMC
(Intercept)                   0.41041 -0.07025  0.93194   1000.0  0.078 .
poly(deltaT, 2)1              3.39203  2.18043  4.80700   1000.0 <0.001 ***
poly(deltaT, 2)2              1.24600 -0.79716  3.56551   1000.0  0.244
clade2multi                   0.37864 -0.36047  1.18762   1000.0  0.284
poly(deltaT, 2)1:clade2multi -2.89770 -5.34509 -0.47838   1000.0  0.018 *
```

poly(deltaT, 2)2:clade2multi 0.66985 -2.37192 3.09101	1000.0 0.614

**SI Table 3.2: Full REML mixed effect model on changes in Log_e_(Δ𝝎_stress_/ Δ𝝎_benign_).**

Random effects:

Groups	Name	Variance	Std.Dev.

source	(Intercept)	0.4429	0.6655

organism	(Intercept)	0.0000	0.0000

Residual	0.5366	0.7325

Number of obs: 94, groups: source, 28; organism, 11

Fixed effects: (contrasts reported against model intercept: Mutational effects measured on growth rate at high temperature stress, for mutations introduced by insertions/deletions)

Fixed effects:

Estimate	Std. Error	t value

Intercept (HighT:ins/del:growth rate)	1.2142	0.3807	3.189

LowT	-1.3192	0.3066	-4.303

Other	-1.1166	0.2510	-4.448

Method:MA	0.5133	0.3652	1.405

Method:mutagenesis	-0.1304	0.4611	-0.283

Measure:reproduction	-0.5927	0.3831	-1.547

Measure:survival	0.1576	0.3569	0.442

Analysis of Deviance Table (Type II Wald F tests with Kenward-Roger df)

F	Df	Df.res	Pr(>F)

stress	11.0072	2	79.339	6.041e-05 ***

method	1.2260	2	17.687	0.3172

measure	2.0718	2	11.788	0.1694

**SI Table 3.3: Estimates of mutational variance across environments**

Mutational variance (ΔV) is typically quantified as the excess variance found among mutation accumulation lines or lineages exposed to mutagenesis (as in this study), relative to variance found among control strains. Alternatively, it is estimated from the among- MA line variance component in an ANOVA. Effects of stress on mutational effects were also analyzed in terms of ΔV, which is expected to follow the same patterns as the changes in the cumulative deleterious fitness effect (mutation load; Δ𝝎). Indeed, when replacing the log-ratio of Δ𝝎 across stressful and benign environments with the log-ratio of ΔV, we found the same qualitative pattern: elevated temperature resulted in more mutational variance relative to the benign conditions (P_MCMC_ =0.02), and this increase was significantly greater than for shifts to cold temperature (P_MCMCM_ = 0.02) or the other forms of stress pooled (P_MCMC_ = 0.05), which did not increase mutational variance (see also Figure 4 in main text).

```
DIC: 238.261

 G-structure:  ~source

       post.mean l-95% CI u-95% CI eff.samp
source     2.928   0.5919    6.036     1000

               ~organism

         post.mean  l-95% CI u-95% CI eff.samp
organism    0.1718 2.059e-07   0.9344     1000

 R-structure:  ~idh(stress):units

            post.mean l-95% CI u-95% CI eff.samp
high.units     7.8552    1.665   16.598     1000
low.units      6.7048    2.177   13.785     1000
other.units    0.8896    0.401    1.496     1450

 Location effects: logratio.V ~ stress

            			post.mean l-95% CI u-95% CI eff.samp pMCMC
Intercept (High temp)   	 2.38102  0.49520  4.26237     1000  0.02 *
Low temp   			-2.91793 -5.17509 -0.67177     1107  0.02 *
other  	    			-1.77037 -3.66188 -0.01586     1000  0.05 *
```

**SI 3.4: Estimating number vs effect of mutations via relationships between mutation load and mutational variance.**

We explicitly modelled increases in mean mutational effects across temperature. Moreover, the analysed studies, including our own, induced a fixed number of mutations to be compared across environments. Nevertheless, environmentally induced changes in mutational effects may in the eyes of natural selection also materialize as increases in the mean number of effectively deleterious (and no longer quasi-neutral) mutations. Assuming that the number of accumulated mutations is approximated by a Poisson process, and that variance in effects of single mutations (coefficients of mutational variation: CV_m_) are constant across environments, we can approximate changes in the number and effect of deleterious mutations across environments by comparing mutation load (Δ𝝎) and mutational variance (ΔV) (Bateman 1959, Mukai 1964). If the observed increase in Δ𝝎 at temperature stress is only due to the number of deleterious mutations, then changes in ΔV are predicted to be proportional to changes in Δ𝝎 across environments. On the other hand, if increases in Δ𝝎 were only due to increases in the average fitness effects, then ΔV ~ Δ𝝎^2^ (Bateman 1959. Mukai 1964). Thus, changes in only the number of mutations should yield: Log(ΔV) ~ Log(Δ𝝎), whereas changes in only mutational effects should yield: Log(ΔV) ~ 2Log(Δ𝝎), and changes in both the number and average effects should yield logarithmic exponents between 1 and 2.

We tested these predictions by regressing Log_e_(ΔV_stress_/ΔV_benign_) on Log_e_(Δ𝝎_stress_/Δ𝝎_benign_) for each stress type by applying Standardized Major Axis regression using the smatr package (Warton et al. 2012) for R. We did not find any significant difference in this relationship between high and low temperature, so we pooled all estimates of temperature stress and compared the relationship to that found for other kinds of stress, resulting in 25 estimates for temperature stress and 38 estimates for other forms of stress. This showed that there was no significant difference in the relationship between the two types of stress (LR = 0.03, df = 1, P = 0.87: slope temperature = 1.73, slope other = 1.68, Fig S3.2c). We also performed an analysis excluding extreme observations, with an arbitrary cut-off set at log-ratios greater than 3, removing five observations. This gave the same qualitative result (difference in the relationship between stressors: LR = 0.49, df = 1, P = 0.49).

For the full dataset, the log-ratio of mutational variance increased with a factor of 1.70 (R^2^ = 0.52, P < 0.001) which 95% CI (1.43-2.03) did not overlap 1, suggesting that increases in Δ𝝎 under stress cannot be explained by increases in the number of deleterious mutations only. For the dataset with extreme observations removed, ΔV increased with Δ𝝎 by a factor of 2.09 (95% CI: 1.67-2.63, R^2^ = 0.31, P < 0.001). Together this suggests that increases in both the number of (conditionally) expressed mutations as well as their average fitness effect are likely to underlie the increase in Δ𝝎 under temperature stress, and thus, that our model provides an accurate representation of the mechanistic basis for temperature-dependent mutational fitness effects

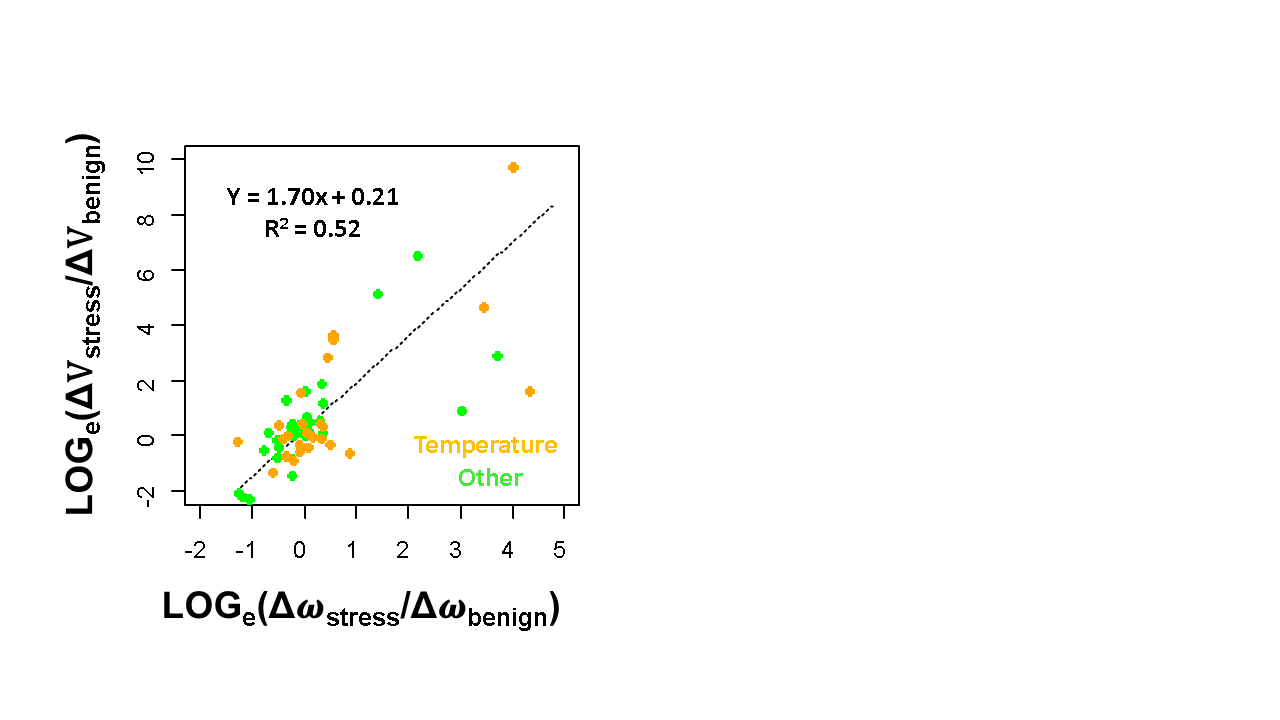

**SI Fig 2.4:** Relationship between the log-ratio of mutational variance (ΔV_stress_/ΔV_benign_) and mutation load (Δ𝝎_stress_/Δ𝝎_benign_) in the stressful and benign environment for the full dataset (temperature stress: orange points, other forms of stress: black points). The hatched line gives the Standardized Major Axis regression slope.

The conclusions above rely on that variation in effects of single mutations (CV_m_) do not change on average across environments (reviewed in Halligan & Keightley 2009). If not, an increase in CV_m_ could contribute to ΔV unequally across environments, and therefore complicate interpretations of the relationship between ΔV and Δ𝝎. Martin & Lenormand (2006) have suggested a method to account for this potential bias by regressing ratios of the Bateman-Mukai estimator of the average mutational effect across stressful and benign environments (s_stress_/s_benign_) on ratios of corresponding variances (ΔV_stress_/ΔV_benign_) (see Martin & Lenormand 2006 for further details). However, since the Bateman-Mukai estimate of s = ΔV/Δ𝝎, this regression violates the assumption of independent measurement error in x and y. There is great uncertainty in estimates of ΔV and s, irrespective of the method used to obtain them (Halligan & Keightley 2009), and regression analysis on this kind of data without accounting for measurement error can lead to very biased estimates and erroneous conclusions (Berger & Postma 2014). Hence, since estimates of measurement error was hard to come by for much of the data, and therefore even harder to correct for, we simply note that there is a potential bias incurred by assuming constant variance in effects of single mutations across environments.

**SI 3. 5: Publication bias.**

Funnel plot showing the distribution of log-ratios against the precision (1/SE) by which they were estimated. Publication bias seems not to have contributed significantly to the reported difference between selection at cold stress (blue) heat stress (red) and other forms of stressors pooled (green). There are many studies reporting non-significant results irrespective of the power of the study. Note that a scaling relationship between the mean log-ratio and its sampling variance is expected since increased mutational effects are expected to result in both increased selection coefficients and mutational variance, the latter being represented in form of differences between genotypes used to assess the sampling variance of selection coefficients. Note also that the three most extreme increases in log-ratios under elevated temperature (red points to the right) occurred in the three experiments imposing the highest increase in temperature, and are thus following expectations.

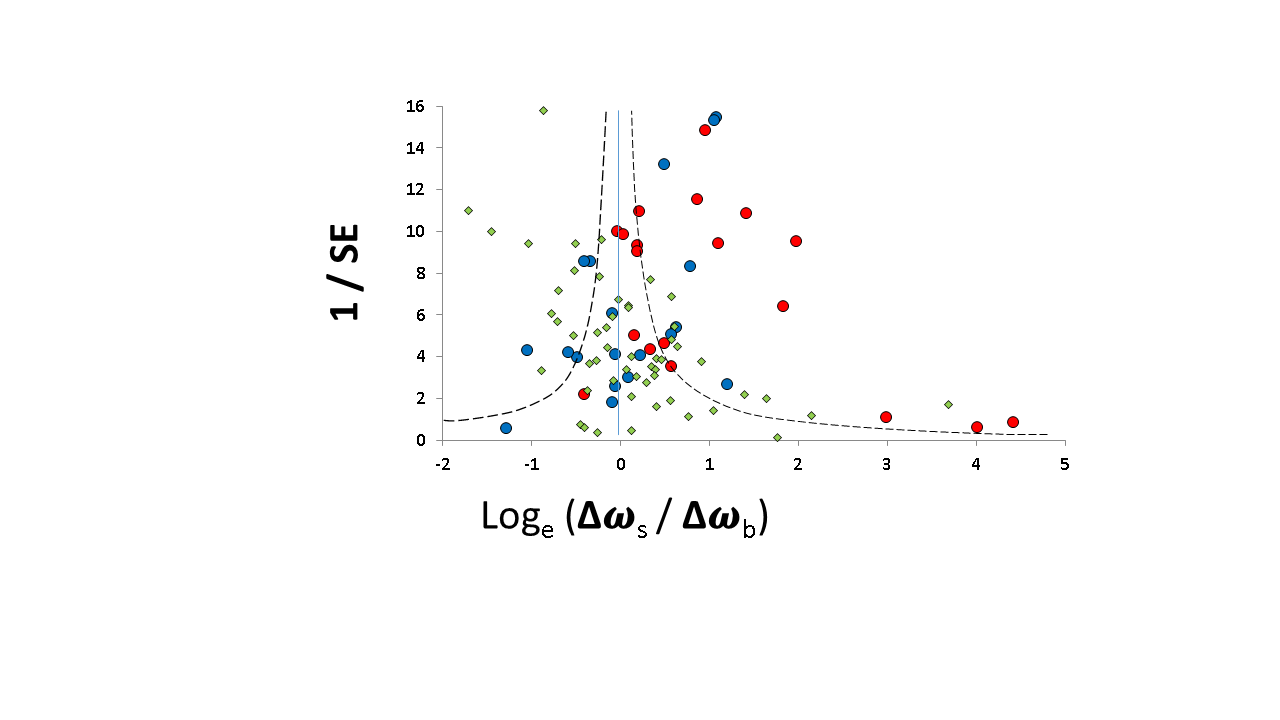
